# Tetracyclines Modify Translation By Targeting Key Human rRNA Substructures

**DOI:** 10.1101/256230

**Authors:** Jonathan D. Mortison, Monica Schenone, Jacob A. Myers, Ziyang Zhang, Linfeng Chen, Christie Ciarlo, Eamon Comer, S. Kundhavai Natchiar, Steven A. Carr, Bruno P. Klaholz, Andrew G. Myers

**Author notes:** Correspondence (A.G.M.).

## Abstract

Apart from their antimicrobial properties, tetracyclines demonstrate clinically validated effects in the amelioration of pathological inflammation and human cancer. Delineation of the target(s) and mechanism(s) responsible for these effects, however, has remained elusive. Here, employing quantitative mass spectrometry-based proteomics, we identified human 80S ribosomes as targets of the tetracyclines Col-3 and doxycycline. We then developed in cell click selective crosslinking with RNA sequence profiling (icCL-Seq) to map binding sites for these tetracyclines on key human rRNA substructures at nucleotide resolution. Importantly, we found that structurally and phenotypically variant tetracycline analogs could chemically discriminate these rRNA binding sites. We also found that tetracyclines both subtly modify human ribosomal translation and selectively activate the cellular integrated stress response (ISR). Together, the data reveal that targeting of specific rRNA substructures, activation of the ISR, and inhibition of translation are correlated with the anti-proliferative properties of tetracyclines in human cancer cell lines.

## INTRODUCTION

Tetracyclines have well-documented antibiotic activity, blocking general protein synthesis in bacteria by binding to the 30S ribosomal subunit, inhibiting binding of tRNAs in the A-site (Brodersen et al., 2000; Pioletti et al., 2001). In addition to this primary mode of action, tetracyclines have also demonstrated remarkably diverse and beneficial “off-target” biological activities in humans, spurring interest in their development as non-antimicrobial therapeutics. A number of natural and semisynthetic tetracyclines have shown favorable anti-inflammatory (Bensman et al., 2012; Cazalis et al., 2009), anti-proliferative (Lokeshwar, 2011), anti-fibrogenic (Xi et al., 2013), neuroprotective (Yong et al., 2004), as well as vaso-and cardio-protective (Fainaru et al., 2008; Tessone et al., 2005) effects in a number of *in vitro* and *in vivo* pre-clinical studies.

Promising results have also emerged from the clinical application of tetracyclines as anti-angiogenic and anti-proliferative agents. In two salient cases, patients with rare diseases, a young male with pulmonary capillary hemangiomatosis (Ginns et al., 2003) and an elderly woman with lymphangioleiomyomatosis (Moses et al., 2006), both showed complete remission after treatment with the tetracycline analog doxycycline. The anti-angiogenic and anti-proliferative activities of doxycycline were hypothesized to be the primary modes of action for inhibiting disease progression and restoring normal pulmonary function in each case.

Numerous clinical trials for tetracyclines in human disease have also shown positive outcomes. The semi-synthetic, non-antimicrobial tetracycline analog Col-3 has shown promise in early clinical trials with patients with AIDS-related Kaposi’s sarcoma (KS) (Dezube et al., 2006) and refractory metastatic cancers (Rudek et al., 2001). Favorable outcomes have also been observed with tetracyclines in indications for rheumatoid arthritis (O’Dell et al., 2006) and osteoarthritis (Brandt et al., 2005), Fragile X syndrome (Leigh et al., 2013), and abdominal aortic aneurysm (Lindeman et al., 2009).

Despite these promising clinical results and a wealth of published scientific research, no relevant target(s) or mechanism(s) have been clearly identified to account for these non-canonical effects of tetracyclines. This imposes a significant barrier to clinical development of tetracyclines in new disease indications and rational drug discovery efforts. It has long been assumed that tetracyclines target bacterial ribosomes selectively and do not bind mammalian ribosomes with any notable functional consequences. Herein, we show that the human ribosome is a relevant biological target of the tetracyclines Col-3 and doxycycline, and these tetracyclines bind a number of important ribosomal RNA (rRNA) substructures that do not overlap with the binding sites of known eukaryotic translation inhibitors (Garreau de Loubresse et al., 2014). Instead, we find that the targeted rRNA substructures comprise eukaryotic ribosomal elements known to be involved in the regulation of translation processes, such as initiation, elongation, and tRNA binding. These sites include helices h16 and h18 on the 40S subunit, as well as H89 and residues at the entrance to the peptidyl exit tunnel on the 60S subunit. Functionally, we show that tetracyclines promote both partial inhibition of ribosomal translation and selective activation of the integrated stress response (ISR). Notably, we found that targeting of human ribosomal RNA structures, inhibition of translation, and activation of the cellular ISR are all strongly correlated to the anti-proliferative effects of both Col-3 and doxycycline in human cancer cell lines.

## RESULTS

### Affinity Isolation and Analysis of Putative Tetracycline Targets

First, to gain an unbiased view of putative proteins that bind tetracyclines, we used affinity isolation in tandem with mass spectrometry (MS)-based quantitative SILAC (stable isotope labeling by amino acids in cell culture) proteomics, a powerful technique that has been used to assign numerous small molecules unambiguously to their cellular binding partners (Ong et al., 2009). We synthesized immobilized derivatives of the tetracycline analogs Col-3 and doxycycline (**Fig. 1a**; **Supplementary Results**, **Supplementary Fig. 1a–b**), for use as affinity probes for SILAC experiments. These tetracycline-based SILAC probes retained modest anti-proliferative activity relative to their respective parent tetracycline scaffold (**Supplementary Fig. 1c**). We incubated our immobilized probes with SILAC-labeled cellular lysates in the presence or absence of soluble Col-3 or doxycycline to competitively bind specific protein targets and thereby diminish their isolation by the affinity matrices, followed by MS analysis of the bound protein fraction (**Fig. 1b**). During the affinity isolation, inclusion of divalent magnesium was essential, for in its absence no proteins were bound (data not shown). This observation is perhaps not surprising, since previously identified targets of the tetracyclines, including the prokaryotic 30S ribosomal subunit (Brodersen et al., 2000; Pioletti et al., 2001), the Tet Repressor (TetR) protein (Hinrichs et al., 1994), and a high affinity tetracycline-binding RNA aptamer (Berens et al., 2001; Xiao et al., 2008), all bind tetracyclines in complex with a magnesium ion. At the same time, 80S ribosomes require magnesium ions for proper folding and assembly (Khatter et al., 2015).

**Figure 1.**
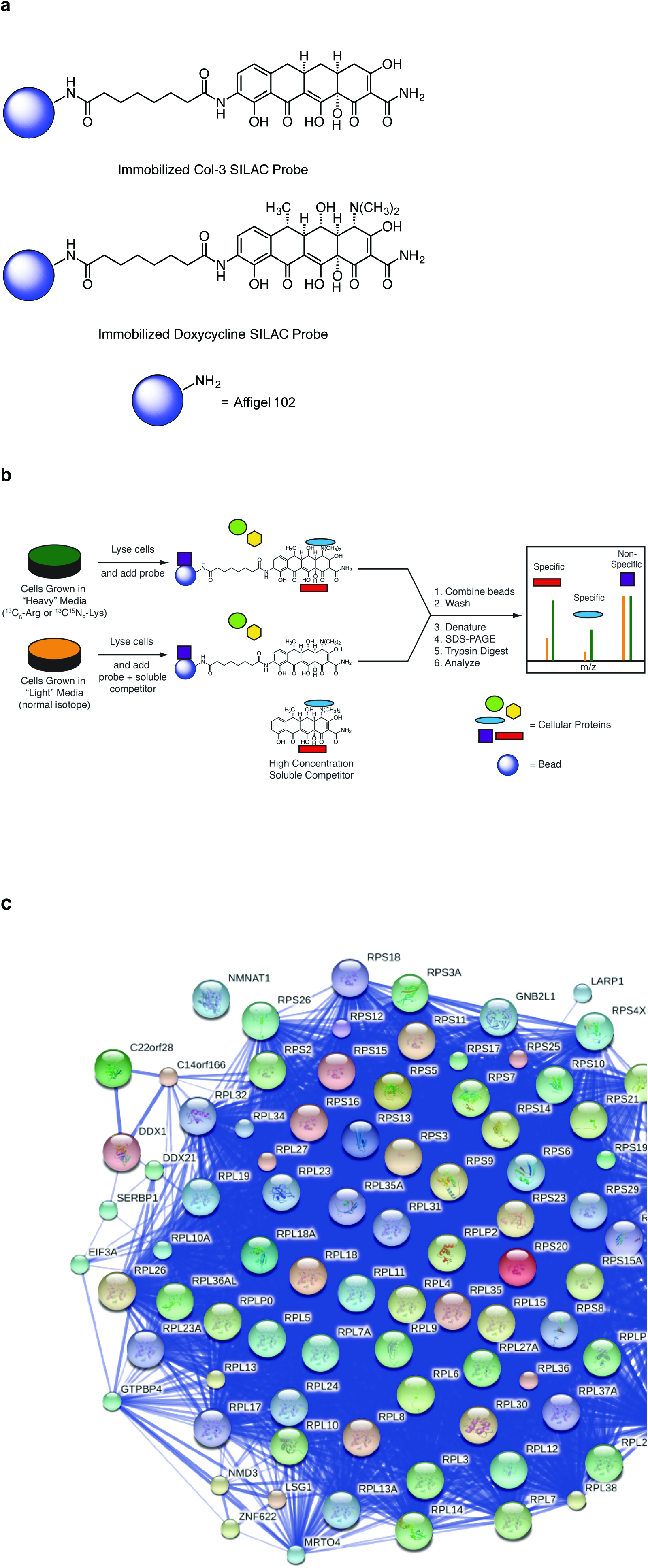
Determination of specific protein binding to tetracyclines. (**a**) Immobilized Col-3 and doxycycline affinity matrices (**b**) Scheme for affinity isolation and tandem MS-based SILAC proteomics. (**c**) A STRING network map of specifically bound proteins (*p* < 0.05) from isolation with tetracycline affinity matrices.

The doxycycline affinity isolation experiment led us to identify 589 proteins, 189 of which were specifically reduced in the presence of free doxycycline (*p* < 0.05) (**Supplementary Fig. 2a**), while the Col-3 affinity isolation experiment revealed 248 proteins, 76 of which were specifically reduced in the presence of free Col-3 (*p* < 0.05) (**Supplementary Fig. 2b**). In the combined datasets, we identified 223 proteins that were bound in both our Col-3 and doxycycline affinity probe experiments, of which 87 proteins were specifically reduced by free drug in the four datasets (*p* < 0.05). Nearly all of the differentially bound proteins from the two datasets were known components of the human 80S ribosomal complex and its closely associated factors (**Figure 1c**; **Supplementary Table 1**). The observation of affinity for the 80S ribosome is not unprecedented, as tetracycline itself has been shown to have weak affinity for isolated, non-translating 80S eukaryotic ribosomes through *in vitro* competition experiments with^3^H radiolabeled tetracycline (Budkevich et al., 2008). This binding interaction, however, was not characterized in detail and was observed to be weaker than tetracycline’s interaction with the bacterial 30S subunit (K_d_ ~30 μm vs 1–2 μm, respectively). At the same time, these previous experiments did not reveal any measureable effects of tetracycline on general eukaryotic translation *in vitro* (specifically, tetracycline was shown not to affect the synthesis of poly-phenylalanine from a poly-uridine transcript).

### In Cell Click Selective Crosslinking with RNA Sequence Profiling Maps Tetracycline Binding Sites on the Human Ribosome

Following the identification of the 80S ribosome as a putative target of Col3 and doxycycline, we next sought to identify the precise binding sites for these tetracyclines within the ribonucleoprotein complex. With the recent development of novel, bifunctional reagents for protein target identification (Li et al., 2013) and *in vivo* mapping of RNA structures (Flynn et al., 2016; Spitale et al., 2015), we developed a hybrid strategy to map ribosomal binding sites of tetracyclines by incorporating dual bioorthogonal handles into tetracycline-based probes. Toward this end, we developed a short synthesis of a versatile linker containing both a photoactive diazirine to enable direct probe crosslinking to the human ribosome and an azide handle to allow selective enrichment of crosslinked biomolecules via copper-free click chemistry (Jewett et al., 2010) (**Supplementary Fig. 3a–b**). This linker was readily incorporated into “active” Col-3 and doxycycline bifunctional probes shown in **Fig. 2a**, which retained equipotent anti-proliferative activity relative to each respective parent tetracycline scaffold (**Fig. 2b**). We also synthesized a non-specific aniline control probe containing our bifunctional linker and, usefully, “inactive” Col-3 and doxycycline probes having no measurable anti-proliferative effects relative to Col-3 and doxycycline, respectively, through incorporation of a slightly modified “inactive” diazirine-azide linker. These probes were then employed for in cell click selective crosslinking with RNA sequence profiling (icCL-Seq) (**Fig. 2c**).

**Figure 2.**
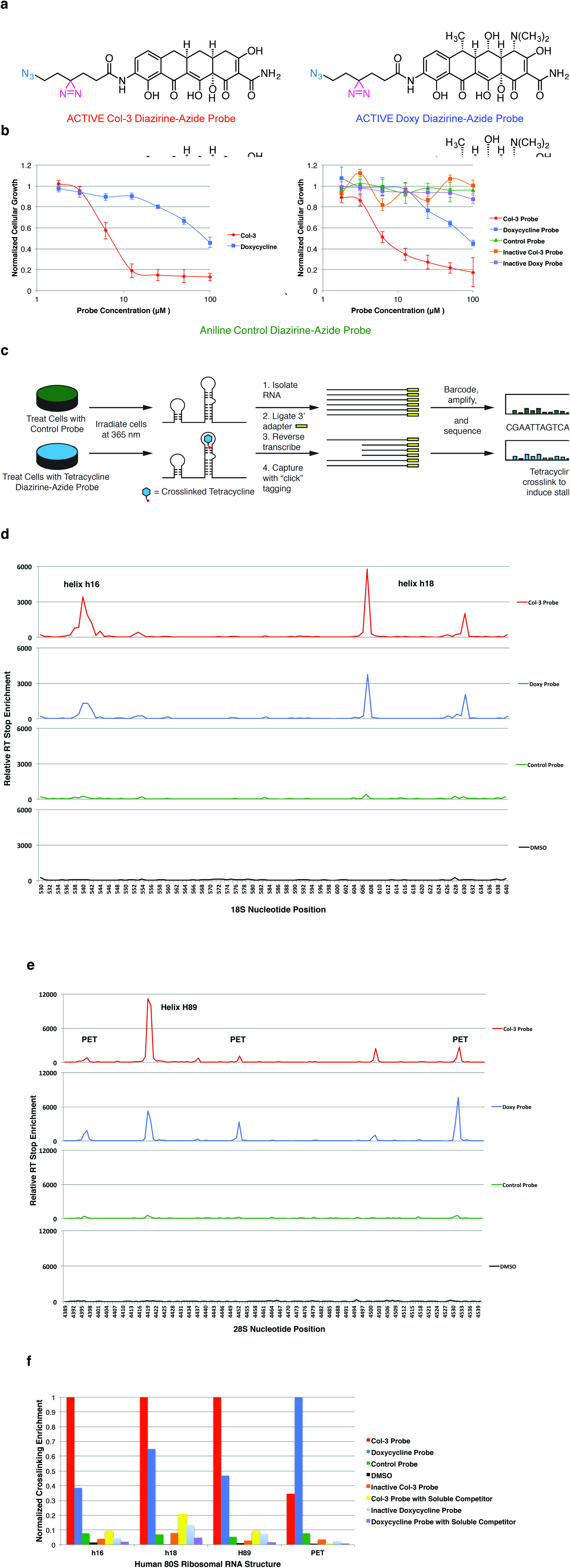
Mapping of tetracycline binding sites on the human ribosome with in cell “click”-selective crosslinking with RNA sequence profiling. (**a**) Bifunctional Col-3 and doxycycline diazirine-azide probes and inactive aniline, Col-3, and doxycycline control probes. The diazirine moiety is highlighted in magenta and the azide is highlighted in teal. (**b**) Scheme for icCL-Seq mapping of binding sites for Col-3 and doxycycline on human ribosomal RNA. (**c**) Reverse transcription stop enrichment map from tetracycline probe crosslinking at helices h16 and h18 on the 18S human ribosomal RNA. (**d**) Reverse transcription stop enrichment map from tetracycline probe crosslinking at helix H89 and the peptidyl exit tunnel on the 28S human ribosomal RNA. (**e**) The relative crosslinking of tetracycline-based bifunctional and control probes at four major binding sites on ribosomal RNAs. Concentration of parent tetracycline soluble competitor was 5X relative to probe for competition studies. PET = peptidyl exit tunnel. (**f**) Growth inhibition curves of A375 cells dosed with Col-3 and doxycycline (left panel) and bifunctional probes (right panel). Data are representative of at least two independent biological experiments.

The active Col-3 and doxycycline probes were each incubated with A375 cells, followed by irradiation at 365 nm to induce photolysis of the diazirine moiety and subsequent crosslinking to adjacent ribosomal components. We also performed crosslinking with our non-specific aniline probe, as well as our inactive Col-3 and doxycycline probes to serve as negative controls (**Fig. 2a**). Pulldown and sequencing of the crosslinked RNAs from our experiments established that active bifunctional tetracycline probes gave the most significant enrichment of reverse transcription (RT) stops at a number of key ribosomal RNA sites (**Fig. 2d** and **Fig. 2e**; **Supplementary Figure 3c**) relative to background and the non-specific and inactive controls. The regions most highly enriched for crosslinking by the active Col-3 and doxycycline probes each had two or more adjacent bases that were crosslinked, which were verified in multiple independent crosslinking and pulldown experiments. These sites corresponded to the apical loop of ribosomal helix h16 (nucleotides 539-542; up to >3,400-fold enrichment), the bulge loops of helix h18 (U607 and U630; up to >5,700-fold enrichment), the terminal loop of helix H89 (U4419–4420; up to >12,000-fold enrichment), and the entrance to the peptidyl exit tunnel (PET) (A4396–4397, U4452, U4530–4531; up to >7,500-fold enrichment). Further, to determine the specificity of these binding and crosslinking events, we also performed in cell crosslinking of our active tetracycline probes in the presence of high concentrations of the respective parent tetracycline drug as a direct binding competitor. It was found that co-incubation of cells with these probes plus parent tetracycline greatly reduced RT stop enrichment at nearly all identified sites (**Fig. 2f**; **Supplementary Figure 3c**), demonstrating successful displacement of each respective probe at ribosomal binding sites by parent Col-3 or doxycycline.

Strikingly, we found little to no crosslinking above background with either of the inactive Col-3 or doxycycline probes at helices h16, h18, H89, or the ribosomal PET (**Fig. 2f**). The active tetracycline-based probes differ from the inactive probes only in the arrangement of the diazirine and azide functional groups, which are compactly displayed in a linear array on the active probes versus a branched array on the inactive probes (**Fig. 2a**). Despite their general structural similarities, only the linearly arrayed active diazirine-azide probe sets have anti-proliferative activity in A375 cells (**Fig. 2b**). These important structure-activity and binding relationships for the tetracyclines highlight that differences in the tetracyclines’ anti-proliferative effects are likely related to differences in their ability to bind at one or more of the ribosomal sites identified in our crosslinking studies.

To further understand the functional implications of tetracyclines binding to the human ribosome, we sought to quantify relative crosslinking differences of active Col-3 and doxycycline probes among the identified binding sites to find potential correlations with the probes’ observed anti-proliferative activity in A375 cells. While both Col-3 and its active bifunctional probe derivate have good anti-proliferative activity and are essentially equipotent (GI_50_ = 9.1 and 7.6 μm, respectively) in A375 cells, doxycycline and its active bifunctional probe derivate have little to no anti-proliferative activity in this cell line at physiologically relevant concentrations (GI_50_ = 92.4 and 80.8 μm, respectively) (**Fig. 2b**). The data showed significantly lower binding and crosslinking enrichment of the active doxycycline probe compared to the active Col-3 probe at h16, h18, and H89, where the relative RT stop enrichments at the most reactive nucleotide in each helix were 27.3, 59.0, and 36.0%, respectively (**Fig. 2f**). By contrast, the active doxycycline probe showed greater crosslinking at the entrance to the PET, where the active Col-3 probe showed only 32.7% relative RT stop enrichment at the most reactive nucleotide. At the same time, we also found that the aniline control, inactive Col-3, and inactive doxycycline probes did not significantly crosslink to any of these ribosomal RNA substructures. Since the active Col-3 probe binds and crosslinks to h16, h18, and H89 to a significantly higher degree than the active doxycycline probe, increased targeting of these regions by Col-3 may explain its stronger anti-proliferative properties relative to doxycycline. Doxycycline, on the other hand, could mediate other effects on the ribosome through stronger binding at the entrance to the peptidyl exit tunnel. Taken together, these findings are suggestive that differential binding of these tetracycline analogs to one or more of these ribosomal RNA sites could lead to differential effects in mammalian cells, especially cellular proliferation.

To gain molecular insights into the nature of the interactions between Col-3 and doxycycline and the ribosome, we performed *in silico* three-dimensional structure analysis of their putative ribosomal binding sites. Based on the recently determined structure of the human 80S ribosome (Khatter et al., 2015), the positions of cross-linked residues can be identified in the structure and visualized. Two sites are located on the 18S rRNA of the 40S ribosomal subunit (**Fig. 3a** and **Fig. 3b**) at the tip of helix h16, as well as on the internal loops of helix h18. These regions are in proximity to the messenger RNA channel and both structures are intimately involved in regulating translation initiation and elongation (Hashem et al., 2013; Martin et al., 2016; Pisarev et al., 2008). Tetracyclines binding to these substructures could thus affect translation initiation or mRNA progression during elongation. Two other sites are on the 28S rRNA of the 60S ribosomal subunit (**Fig. 3c** and **Fig. 3d**) at the apical loop of helix H89 and at the entry of the PET, which are both important sites for elongation and egress of the nascent peptide during translation. It is interesting to note that all of these binding sites are distinct from vestigial sites of bacterial ribosomal binding of tetracyclines (Brodersen et al., 2000; Pioletti et al., 2001), and are also distinct from known binding sites of eukaryotic ribosomal inhibitors (Garreau de Loubresse et al., 2014), and thus may represent new targets on the human ribosome for small molecules. In addition, closely associated magnesium ions can be observed in three of the four binding sites, which could serve to mediate the tetracyclines’ interaction with the ribosomal RNA. While the tetracycline ligands cannot yet be modeled into these sites with reasonable precision, this analysis illustrates clear potential binding pockets as supported by our crosslinking data (magenta spheres in **Fig. 3a–d**). Interestingly, none of these binding sites are proximal to ribosomal structural proteins (i.e. drug interactions appear to be mediated solely via RNA elements of the human ribosome).

**Figure 3.**
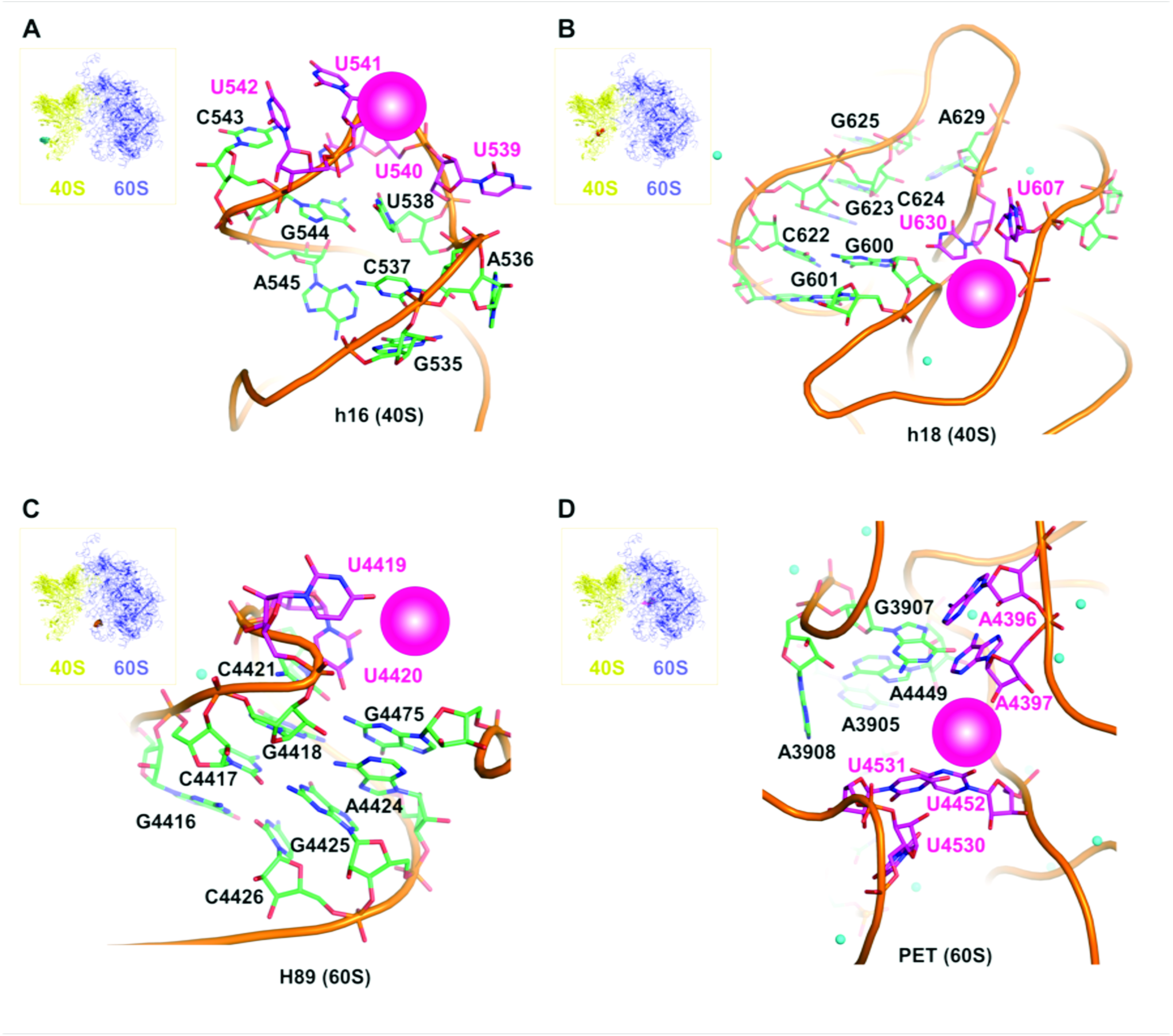
Modeled structural representations of putative binding sites for the tetracyclines Col-3 and doxycycline. Binding sites on the small, 40S subunit at (**a**) h16 and (**b**) h18. Binding sites on the large, 60S subunit at (**c**) H89 (**d**) peptidyl exit tunnel (PET). Crosslinked bases are highlighted in magenta and magenta spheres represent potential binding pockets for Col-3 and doxycycline at each site. Small green spheres are divalent magnesium ions.

### Tetracyclines Activate the Integrated Stress Pathway Selectively

To learn which mammalian cellular pathways are affected by tetracyclines, we performed RNA-Seq to identify changes in gene expression in cells upon treatment with Col-3 or doxycycline. After a 6-hour treatment with these two analogs in A375 cells, we observed an enriched gene signature corresponding to activation of the cellular unfolded protein response (UPR) and ER stress pathways in cells treated with 10 μm Col-3. This enrichment was absent in cells treated with 10 μm doxycycline. Interestingly, this gene signature was most highly enriched for targets of activating transcription factor 4 (ATF4), such as *CHOP, GADD34, ATF3*, and *SESN2* (**Fig. 4a**; **Supplementary Table 2**), corresponding to specific activation of the integrated stress response (ISR) pathway, a specialized branch of the UPR (Ron and Walter, 2007).

**Figure 4.**
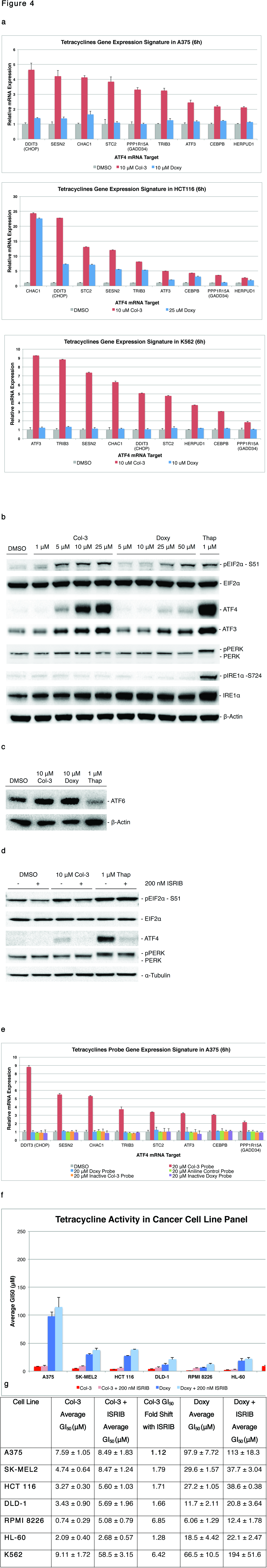
Activation of the ISR by tetracyclines in A375 cells. (**a**) Col-3 but not doxycycline activate ATF4 target genes. (**b**) Tetracyclines induce dose-dependent phosphorylation of EIF2α and ATF4 expression at 6 h, but not phosphorylation of PERK or IRE1α. Thapsigargin (Thap), was used as a positive control for induction of ER stress. (**c**) Col-3 and doxycycline do not stimulate cleavage and activation of ATF6 at 6 h. (**d**) Co-treatment of A375 cells with ISR inhibitor blocks Col-3-mediated activation of ATF4 (**e**) The Col-3-based bifunctional probe induces expression of ATF4 gene targets in A375, while the gene signature is not activated by the doxycycline-based probe, the aniline control probe, or the inactive tetracycline-based bifunctional probes. (**f**) Average growth inhibition (GI_50_) of A375, SK-MEL2, HCT 116, DLD-1, RPMI 8226, HL-60, and K562 cells co-dosed with tetracyclines and ISRIB. (**g**) Tabular GI_50_ values for A375, SK-MEL2, HCT 116, DLD-1, RPMI 8226, HL-60, and K562 cells co-dosed with tetracyclines and ISRIB Data are represented as mean ± SEM. Data are representative of experiments with at least three independent experiments.

Stimulation of the ISR occurs through upstream signaling of cell-stress sensor kinases, which all converge on a single substrate and cellular event, the phosphorylation of the α-subunit of eukaryotic translation initiation factor 2 (*p*-eIF2α) at serine-51 (Ron and Walter, 2007). We looked at the effects of Col-3 and doxycycline on signaling events upstream of ATF4 and found that while Col-3 induced robust, dose-dependent eIF2α phosphorylation, doxycycline did not induce these effects at physiologically relevant concentrations (**Fig. 4b**). Instead, only low levels of eIF2α phosphorylation and ATF4 expression were observed in A375 cells treated at a high dose (50 μm) of doxycycline. We also found that Col-3 induces phosphorylation of eIF2α independently of the general upstream regulator of ER stress, protein kinase R-like endoplasmic reticulum kinase (PERK). As shown in **Fig. 4b** and **Fig. 4c**, activation of the ISR by Col-3 was isolated from the initiation of the larger ER stress program, as Col-3 did not activate the other primary ER stress signaling nodes (ATF6 and IRE1α), while the known general ER stress activator thapsigargin (an inhibitor of sarcoplasmic calcium-adenosine triphosphatase) activated all three ER stress branches.

Further evidence that tetracyclines activate ATF4 in an eIF2α-dependent manner was obtained by co-incubating cells with tetracyclines and an ISR inhibitor (ISRIB), which renders cells insensitive to eIF2α phosphorylation through dimerization of the eIF2β subunit of the eIF2 complex(Sidrauski et al., 2015). Co-incubation of A375 cells with Col-3 and ISRIB strongly attenuates ATF4 expression while phosphorylation of eIF2α is largely unaffected (**Fig. 4d**). Additionally, we found that ATF4 induction by tetracyclines strongly correlated with their anti-proliferative activities. At concentrations near its GI_50_ value (~1–10 μm), Col-3 stimulated the expression of an ATF4 target-enriched gene signature in different cancer cell lines, while doxycycline induced expression of ATF4 and its target genes only in doxycycline-sensitive cell lines (e.g., HCT 116 colon carcinoma cells) (**Fig. 4a**). Consistent with these results, only the active Col-3 probe activated the ISR gene signature in A375 cells, while ISR activation was absent with the active doxycycline probe, aniline control probe, and the inactive Col-3 and doxycycline probes (**Fig. 4e**).

Co-administration of ISRIB with tetracyclines might be expected to confer cellular resistance to the anti-proliferative effects observed upon administration of tetracyclines alone; however, in practice we found that conference of resistance by ISRIB co-administration was highly dependent on cancer cell type. While co-treatment of most cell lines with tetracyclines and ISRIB did not result in significant observed resistance, some cell lines (RPMI 8226 and K562) showed strong resistance (**Fig. 4f**) upon co-treatment with tetracyclines and ISRIB. These data suggest that while ISR induction may play a role in the anti-proliferative effects of the tetracyclines, they do not fully explain these effects across all cancer cell types. Accordingly, ISR induction by Col-3 and doxycycline may play an ancillary role in reducing proliferation in some cellular contexts and a major role in other cellular contexts. At the same time, these findings suggest that Col-3 and doxycycline likely modulate additional pathways apart from the ISR that serve to diminish cellular proliferation.

### Tetracyclines Can Inhibit Human Translation Independent of ISR Activation

Following identification of the human ribosome as a target of Col-3 and doxycycline, we sought to define the consequences of their ribosomal binding by studying the effects of these tetracycline analogs on global cellular translation. By using *O*-propargyl puromycin (OPP), which like puromycin is taken up by cells and incorporated into nascent peptides within ribosomes, we could track *de novo* protein synthesis in live cells (Liu et al., 2012) by detecting levels of OPP incorporation following tetracycline treatment. This is accomplished via fluorescent tagging of OPP-labeled nascent peptides in cells using click chemistry (Kolb et al., 2001), followed by quantitation of the tagged peptides by flow cytometry.

We treated A375 cells for 3 hours with either Col-3 or doxycycline at concentrations of both 10 and 25 μm, as well as with either 10 μm cycloheximide as a positive control or vehicle alone as a negative control. We then pulse labeled cellular nascent peptides with OPP for 1 h, followed by tagging of OPP-labeled peptides with Alexa Fluor 488 azide and analysis by flow cytometry. Surprisingly, we found that treatment of A375 cells with either 10 or 25 μm Col-3 resulted in a small, but discernible decrease in cellular protein synthesis (~26 and 36% inhibition, respectively); however, translation inhibition was not nearly as pronounced as that observed with the general eukaryotic translation inhibitor cycloheximide (>80% inhibition) (**Fig. 5a** and **Fig. 5b**). With doxycycline, there was no observable change in cellular translation at a concentration of 10 μm doxycycline and only a slight decrease in translation at a concentration of 25 μm doxycycline (~16% inhibition). These findings provide strong evidence that Col-3 and doxycycline affect translation in a dose dependent manner through direct binding to the human ribosome. At the same time, the differential effects on translation inhibition by Col-3 relative to doxycycline may underlie the disparate anti-proliferative effects of these two analogs.

**Figure 5.**
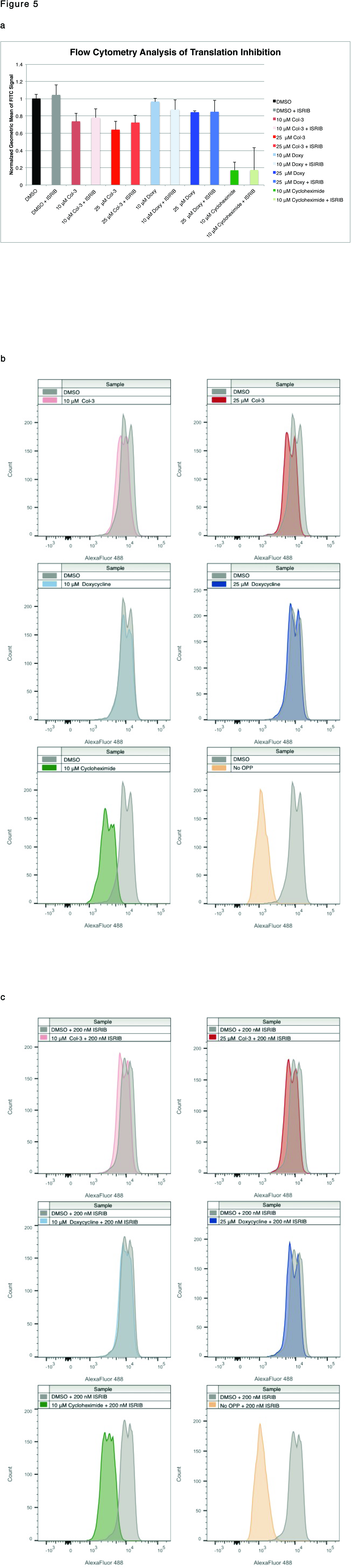
OPP-flow cytometry analysis shows tetracyclines inhibit translation independent of activation of the ISR. (**a**) Relative OPP incorporation in nascent peptides from tetracycline-treated A375 cells. Data are represented as mean ± SEM (**b**) Histogram plots for translation analysis in A375 cells dosed with tetracyclines at 3 h (**c**) Histogram plots for translation analysis in A375 cells co-dosed with tetracyclines plus 200 nM ISRIB at 3 h. Data represents experiments with a minimum of three independent replicates.

Finally, since activation of the ISR results in changes to mammalian translation through reduced abundance of functional initiator methionine tRNA complexes (eIF2–GTP–tRNAi^Met^), we next sought to learn whether tetracyclines have discernible effects on translation apart from ISR induction. We co-dosed A375 cells with either Col-3 or doxycycline plus ISRIB and found that ISRIB co-treatment did not restore global protein synthesis to levels seen in control-treated cells. Rather, treatment with tetracyclines plus ISRIB resulted in comparable levels of translation inhibition relative to the levels observed upon treatment with tetracyclines alone (**Fig. 5a** and **Fig. 5c**). As expected, ISRIB co-treatment did not affect cycloheximide’s ability to block translation. These results suggest that protein synthesis changes induced by tetracyclines are not primarily driven by ISR activation, but instead result from direct inhibition of the human ribosome. Since translation inhibition and ribosomal stalling have been shown to promote eIF2α phosphorylation (**Supplementary Figure 4**) and activation the ISR (Ishimura et al., 2016), tetracycline-induced ribosome stalling may provide the primary upstream event that leads to initiation of the ISR by tetracyclines.

## DISCUSSION

The effects of tetracyclines on human ribosomal function described herein highlight a novel paradigm for these drugs and dispel notions that antibiotics such as the tetracyclines, which are well recognized to target bacterial ribosomes, do not also have functional consequences on their eukaryotic counterparts. To our knowledge, this represents the first direct evidence that tetracyclines have observable effects on human cytosolic ribosomes, including direct inhibition of translation and subsequent downstream activation of the integrated stress response. In this work, we also demonstrate a method for mapping small molecule-RNA interactions in live cells through icCL-Seq. Through conjugation of a versatile bifunctional linker containing a photoreactive diazirine and a “click”-enabled azide handle to Col-3 and doxycycline, we generated probes capable of selective crosslinking and enrichment of ribosomal RNA binding sites for these tetracycline analogs at nucleotide resolution. We envision this strategy could be generally applicable to the discovery of other novel RNA targets of small molecules.

Our work reports that tetracyclines bind divergent human ribosomal substructures from those targeted by other known small molecule eukaryotic translation inhibitors (Garreau de Loubresse et al., 2014), revealing new sites on the human ribosome for potential targeting. Conceptually, this finding mirrors the identification of multiple binding sites (2–6 sites of varying occupancy) reported for tetracycline on the prokaryotic ribosome in independent crystal structures and biochemical data (Brodersen et al., 2000; Pioletti et al., 2001). We have identified at least four high-confidence binding sites for the tetracyclines Col-3 and doxycycline on the human ribosome, three of which show good correlation between tetracycline binding and tetracycline anti-proliferative activity. While it is still speculative at this time, differential binding of Col-3 relative to doxycycline at helices h16, h18, and H89, as well as a lack of binding of our inactive tetracycline probes at these three sites, implicates these sites as mediators of the tetracyclines’ anti-proliferative properties. Continued work will be necessary to better understand how the tetracyclines’ binding/avidity for these (or still yet to be discovered) sites could affect ribosome function, and we envision that it may be possible to synthesize tetracycline analogs that can uniquely target single ribosomal binding pockets with high selectivity.

Targeting translation on human cytosolic ribosomes is an emerging strategy for therapeutic intervention in human diseases with dysregulated protein synthesis such as cancer (Bhat et al., 2015; McClary et al., 2017; Myasnikov et al., 2016) with the eukaryotic translation inhibitor homoharringtonine (omacetaxine) recently entering the clinic for treatment of chronic myelogenous leukemia that is refractory to tyrosine kinase inhibition therapy (Gandhi et al., 2014). This work adds tetracyclines to the growing class of small molecules with therapeutic potential in targeting the human ribosomal translational machinery. In this article, we have shown that the tetracyclines Col-3 and doxycycline induce dose-dependent reductions in human ribosomal translation that strongly correlate with their cellular antiproliferative activities and are distinct from their effects on the cellular ISR. These effects on translation were not as profound as those observed with the translation inhibitor cycloheximide. Instead, Col-3 and doxycycline only partially inhibit human ribosomal translation at clinically relevant concentrations (Syed et al., 2004), representing a potential therapeutic window for tetracycline analogs as antiproliferative agents against cancer cells through targeting of protein synthesis.

Further studies are required to explore the tetracyclines’ mechanism of action on the ribosome in detail, especially with regard to their perturbation of protein translation and activation of the ISR. Of particular interest are any selective effects tetracyclines may have on the human translational program and how each of the unique binding sites of tetracyclines on the human ribosome may influence those effects. Previous studies have demonstrated that both the ribosomal mRNA tunnel and nascent peptide tunnel are susceptible to modification in ways that induce translational stalling or slowing of particular transcripts. Gupta and co-workers have shown that ribosomal modifications in bacteria, such as methylation by *erm* resistance genes, can induce stalling of specific transcripts that in turn reduces bacterial fitness (Gupta et al., 2013). At the same time, macrolide antibiotics have been shown to selectively influence translation in bacteria both through perturbation of the nascent peptide exit tunnel (Kannan et al., 2012), as well as through allosteric modification of the catalytic peptidyl transferase center (Sothiselvam et al., 2014). In mammalian cells, both of the tetracycline-targeted helices h16 and h18 are directly involved in dynamic translation initiation and elongation processes. Specifically, h16 binds a number of regulatory proteins such as the helicase DHX29 (Hashem et al., 2013; Pisareva et al., 2008), which unwinds mRNAs with highly structured 5’-untranslated regions, and can also directly base pair with RNAs entering the mRNA channel (Martin et al., 2016). Helix h18 directly interacts with incoming messenger RNAs (Pisarev et al., 2008) and undergoes many conformational changes during initiation (Passmore et al., 2007). In light of these findings, it could be envisioned that tetracyclines inhibit human ribosomal translation by perturbing local rRNA structures near the mRNA or nascent peptide tunnels, or by modifying the association or organization of other higher order translational regulatory elements on the human ribosome.

## SIGNIFICANCE

Tetracyclines have demonstrated clinically relevant activity in the inhibition of cancer proliferation; however, to date, the relevant targets responsible for these anti-proliferative activities have not been clearly identified. We have employed a multi-disciplinary approach to identify the human ribosome as a target of the tetracyclines Col-3 and doxycycline, map the ribosomal binding sites for these tetracyclines analogs on human rRNA substructures, and define the tetracyclines’ mechanism of action as a partial human ribosomal translation inhibitor. This work lays the foundation for future structural and mechanistic experiments examining tetracycline function on the human ribosome while also paving the way for rational clinical development of superior tetracyclines for human diseases such as cancer.

## STAR METHODS

Methods, including statements of data availability and any associated accession codes and references, are available in the online version of the paper.

### Reagents and Cell Culture Details

A375, HCT 116, K562, and HeLa cell lines were obtained from ATCC (CRL-1619™, CCL-247™, CCL-243™, and CCL-2™, respectively). A375 and HeLa cells were cultured in Dulbecco’s Modified Eagle Medium (DMEM) supplemented with 10% fetal bovine serum. HCT 116 cells were cultured in McCoy’s 5a (Iwakata & Grace Modification) media supplemented with 10% fetal bovine serum. K562 cells were cultured in Iscove’s Modification of DMEM (IMDM) media supplemented with 10% fetal bovine serum. All cell cultures were maintained in a 5% CO_2_ incubator at 37 °C. Fetal bovine serum was obtained from Seradigm (Cat. # 1400-100, Lot # 081A11R)

Col-3 was prepared by semi-synthesis from sancycline (*vide infra*), and 2-oxepane-1,5-dione was prepared as previously described(Latere et al., 2002). Doxycycline was purchased from Frontier Chemicals (Cat. # D10056). Sterile DMSO for reagent preparation was obtained from Sigma-Aldrich (Cat. # D2650).

Reactions were performed under ambient atmospheric conditions, unless otherwise noted. Air-and moisture-sensitive reactions, where denoted, were performed in flame-dried glassware fitted with rubber septa under a positive pressure of argon. Air- and moisture-sensitive liquids were transferred via syringe or stainless steel cannula. Solutions were concentrated by rotary evaporation below 35 °C. Analytical TLC was performed using glass plates pre-coated with silica gel (0.25 mm, 60 Å pore size, 230–400 mesh, Merck KGA) impregnated with a fluorescent indicator (254 nm). TLC plates were visualized by exposure to ultraviolet light (UV), followed by staining by submersion in an aqueous, basic solution of potassium permanganate, followed by brief heating on a hot plate. Commercial solvents and reagents were used as received with the following exceptions for anhydrous solvents. Methanol was distilled from calcium hydride under a nitrogen atmosphere at 760 mmHg. Tetrahydrofuran, diethyl ether, dichloromethane, acetonitrile, and dimethyl formamide were purified by passage through Al_2_O_3_ under argon by the method of Pangborn et al. 1,4-cyclohexandione and *p*-toluenesulfonyl chloride were obtained from TCI. Hydroxylamine-*O*-sulfonic acid, *meta*-chloroperoxybenzoic acid, 7N ammonia in methanol, and triethylamine were obtained from Sigma-Aldrich. Flash chromatography was carried out on a Teledyne ISCO CombiFlash purification system with pre-loaded silica gel columns.

Warning: Exercise caution when handling diazirine-azide containing linker compounds as they are potentially high-energy compounds. Working in < 1 g quantities and storage below −20 °C is strongly advised.

### Nuclear Magnetic Resonance

Proton nuclear magnetic resonance spectra (^1^H NMR) spectra were recorded on Varian INOVA 600 (600 MHz), Varian INOVA 500 (500 MHz) or Varian 400 (400 MHz) NMR spectrometers at 23 °C. Proton chemical shifts are expressed in parts per million (ppm, δ scale) and are referenced to residual protium in the NMR solvent (CHCl_3_: δ 7.26, HDO: δ 4.79, CD_2_HOD: δ 4.87, THF-d8: δ 3.58). Data are represented as follows: chemical shift, multiplicity (s = singlet, d = doublet, t = triplet, q = quartet, dd = doublet of doublets, dt = doublet of triplets, td = triplet of doublets, dq= doublet of quartets, dquint = doublet of quintets, sxt = sextet, m = multiplet, br = broad), integration, and coupling constant (J) in Hertz (Hz).

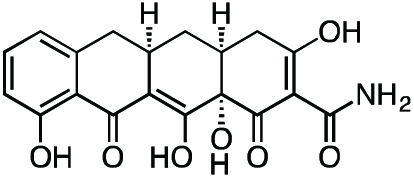

#### Col-3

Methyl iodide (3.27 mL, 52.4 mmol) was added to a stirring suspension of sancycline (0.434 g, 1.047 mmol) in 10 mL THF at 23 °C. The flask was fitted with a reflux condenser, and the reaction mixture was heated at 45 °C for 19 h. The reaction mixture was diluted with 20 mL hexanes and cooled on ice to 4 °C to produce a yellow precipitate. The crude solid was collected by filtration and dried under vacuum. The crude solid was then suspended in 10 mL 1:1 acetic acid/water, followed by the addition of zinc dust (0.343 g, 5.24 mmol). After 45 min, the unreacted zinc was filtered off and the filter washed with excess dichloromethane. The filtrate was concentrated to a dark, yellow-brown solid. The crude solid was dissolved in 30 mL of 20% water/acetonitrile + 0.1% TFA and syringe filtered through a 0.2 micron PTFE filter. The crude filtrate was purified by preparatory RP-HPLC on an Agilent Prep-C18, 10 μm, 21.2 x 250 mm column (20 to 80% CH3CN/H2O + 0.1% TFA, 45 min, 20 mL/min) over 3 runs to provide Col-3 (0.112 g, 28.8%) as its TFA salt as a pale, tan-yellow solid. ^1^H NMR (CD_3_OD, 500 MHz) δ 7.39 (dd, *J* = 8.4, 7.4 Hz, 1H), 6.76 (dd, *J* = 26.8, 7.9 Hz, 1H), 3.27 (dd, *J* = 18.5, 5.6 Hz, 1H), 2.91 (ddd, *J* = 19.4, 10.3, 5.0 Hz, 1H), 2.83 (dd, *J* = 15.2, 4.3 Hz, 1H), 2.53 (t, *J* = 14.7 Hz, 1H), 2.47 (ddd, *J* = 10.5, 5.7, 2.6 Hz, 1H), 2.40 (d, *J* = 18.4 Hz, 1H), 2.05 (ddd, *J* = 13.7, 5.4, 2.8 Hz, 1H), 1.58 (td, *J* = 13.4, 10.8 Hz, 1H). HRMS ESI+ (m/z): 372.1064 (Predicted [M+H]^+^ for C_17_H_17_NO_7_ is 372.1078).

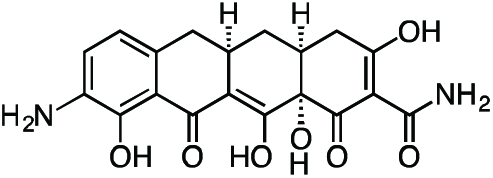

#### 9-Amino-Col-3

Col-3 (0.305 g, 0.821 mmol) was dissolved in 8 mL sulfuric acid and 2 mL methanol, followed by addition of potassium nitrate (0.108 g, 1.06 mmol) at 4 °C. After 2.5 h, the reaction mixture was slowly poured into 50 mL ice-cold (4 °C) water, producing a brown/yellow precipitate. Dichloromethane (100 mL) was added to solubilize the precipitate and the biphasic solution was filtered through Celite, washing the filter with excess dichloromethane. The layers were separated, followed by extraction of the aqueous portion with 4 x 50 mL dichloromethane. Ethyl acetate (100 mL) was added to solubilize the residual precipitate on the Celite filter, followed by washing of the filter with excess ethyl acetate. The layers were separated, followed by extraction of the aqueous portion with 3 x 50 mL ethyl acetate. The combined organic fractions were dried over magnesium sulfate, filtered and concentrated to give a crude mixture of the C7-and C9-nitration products as a yellow-orange solid.

The crude material was re-suspended in 8 mL 1:1 dioxane methanol followed by addition of 20% wt Pearlman’s catalyst (0.115 g, 0.164 mmol) to give a black suspension. The flask was fitted with 3-way stopcock and evacuated and backfilled three times with argon. The stopcock was fitted with a hydrogen balloon, and the flask was then evacuated and backfilled three times with hydrogen. Hydrogen was bubbled into the reaction solution via a long needle extending from the hydrogen balloon. The reaction mixture was stirred vigorously at 23 °C, and the solution phase of the reaction mixture turned a dark red-orange color over 30 min. After 30 min, the residual palladium was filtered off from the reaction mixture through a Celite plug and the crude filtrate was concentrated. The crude residue was re-suspended in 30 mL 50% CH3CN/H2O + 0.1% TFA and filtered through a cotton plug. The filtrate was syringe filtered through a 0.2 micron PTFE filter and purified by preparatory RP-HPLC on an Agilent Prep-C18, 10 μm, 21.2 x 250 mm column (5 to 40% CH3CN/H2O + 0.1% TFA, 45 min, 20 mL/min) over three runs. The C7-amino minor isomer eluted first. The desired C9-amino-Col-3 major isomer eluted second and was isolated as its TFA salt as a yellow-brown solid (0.082 g, 25.8%). ^1^H NMR (CD_3_OD, 500 MHz) δ 7.41 (d, *J* = 8.0 Hz, 1H), 6.85 (d, *J* = 8.1 Hz, 1H), 3.25 (dd, *J* = 18.0, 6.0 Hz, 1H), 2.98 – 2.89 (m, 1H), 2.86 (dd, *J* = 15.4, 4.4 Hz, 1H), 2.55 (t, *J* = 14.9 Hz, 1H), 2.51- 2.43 (m, 1H), 2.39 (d, *J* = 18.5 Hz, 1H), 2.05 (ddd, *J* = 13.5, 5.2, 2.6 Hz, 1H), 1.59 (td, *J* = 13.4, 10.9 Hz, 1H). HRMS ESI+ (m/z): 387.1187 (Predicted [M+H]^+^ for C_19_H_19_N_2_O_7_ is 387.1181).

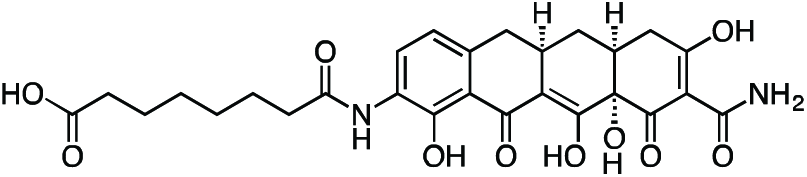

#### Col-3 SILAC Probe

A flask was charged with 9-amino-Col-3 (0.035 g, 0.091 mmol), HATU (0.052 g, 0.136 mmol), and suberic acid (0.079 g, 0.453 mmol). The reaction components were dissolved with 1 mL DMF, followed by addition of triethylamine (25 μL) to give a dark orange solution. The reaction mixture was then stirred for 90 min at 23 °C. The crude reaction mixture was diluted to 10 mL with 50% CH3CN/H2O + 0.1% TFA, syringe filtered and purified by prep RP-HPLC on an Agilent Prep-C18, 10 μm, 21.2 x 250 mm column (20 to 60% CH3CN/H2O + 0.1% TFA, 45 min, 20 mL/min). The Col-3 SILAC probe carboxylic acid was isolated as its TFA salt a yellow solid (16.4 mg, 33.4%). ^1^H NMR (CD_3_OD, 500 MHz) δ 8.04 (d, *J* = 8.1 Hz, 1H), 6.73 (d, *J* = 8.5 Hz, 1H), 3.26 (dd, *J* = 18.1, 5.5 Hz, 1H), 2.96-2.86 (m, 1H), 2.81 (dd, *J* = 15.3, 4.3 Hz, 1H), 2.55-2.44 (m, 2H), 2.46 (t, *J* = 7.4 Hz, 2H), 2.40 (d, *J* = 18.5 Hz, 1H), 2.31 (t, *J* = 7.4 Hz, 2H), 2.04 (ddd, *J* = 13.6, 5.3, 2.8 Hz, 1H), 1.73 (p, *J* = 7.5 Hz, 2H), 1.65 (p, *J* = 7.5 Hz, 2H), 1.59 (td, *J* = 13.4, 10.7 Hz, 1H), 1.49-1.40 (m, 4H). HRMS ESI+ (m/z): 543.1961 (Predicted [M+H]^+^ for C_27_H_30_N_2_O_10_ is 543.1973).

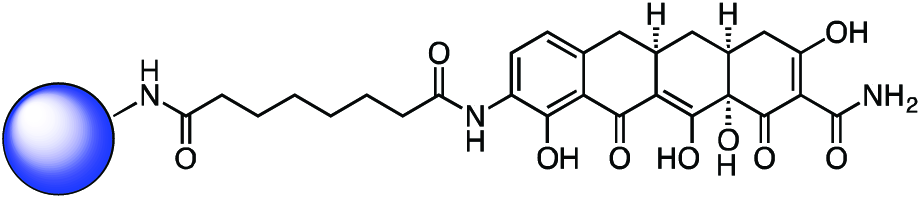

#### Immobilized Col-3 SILAC Probe

To a solution of Col-3 SILAC probe carboxylic acid (**4**) (4.5 mg, 8.29 μmol) and 1-hydroxypyrrolidine-2,5-dione (1.91 mg, 0.017 mmol) in DMSO (800 μL) was added EDC (3.18 mg, 0.017 mmol). After 3h, additional 1-hydroxypyrrolidine-2,5-dione (1.91 mg, 0.017 mmol) and EDC (3.18 mg, 0.017 mmol) were added. After 4h LC-MS analysis showed 84% activation of the starting material corresponding to 6.96 μmol of activated material.

Affi-Gel 102 beads (12 μmol) were washed with 5 x anhydrous DMSO. The beads were then suspended in anhydrous DMSO (500 μL). The activated solution (1.5 μmol) was added to the bead suspension based on the 12.5% loading level and to this was added triethylamine (8.36 μL, 60.0 μmol). The reaction mixture was shaken at RT for 1hr at which point LCMS analysis showed complete loss of signal for the activated material. The beads were washed with DMSO and PBS, followed by final suspension in PBS (1.5 mL) for storage of the immobilized Col-3 SILAC probe.

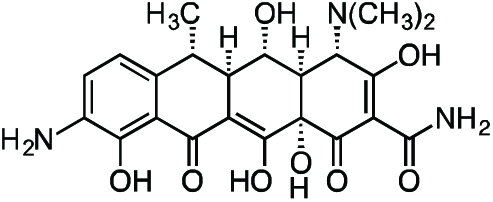

#### 9-Amino-Doxycycline

Nitration of doxycycline was carried out as previously described(Barden et al., 1994): Doxycycline monohydrate (1.04 g, 2.25 mmol) was dissolved in 4 mL concentrated sulfuric acid, followed by the addition of sodium nitrate (0.298 g, 3.51 mmol) at 4 °C. After 3 h, the reaction mixture was added dropwise to a stirring solution of cold ether (150 mL) at 4 °C to produce a yellow precipitate. The precipitate was washed with 3 x 100 mL cold ether and collected by vacuum filtration as a dark yellow solid.

The crude mixture was re-suspended in methanol (40 mL) followed by addition of 20% wt Pearlman’s catalyst (0.315 g, 0.45 mmol) to give a black suspension. The flask was fitted with 3-way stopcock and evacuated and backfilled three times with argon. The stopcock was fitted with a hydrogen balloon, and the flask was then evacuated and backfilled three times with hydrogen. Hydrogen was bubbled into the reaction solution via a long needle extending from the hydrogen balloon. The reaction mixture was stirred vigorously at 23 °C, and the solution phase of the reaction mixture turned a dark red-orange color over 1 h. The crude filtrate was concentrated, re-suspended in 80 mL H2O + 0.1% TFA, syringe filtered through a 0.2 micron PTFE filter, and purified by preparatory RP-HPLC on a Waters SunFire Prep-C18, 10 μm, 21.2 x 250 mm column (0 to 20% CH3CN/H2O + 0.1% TFA, 45 min, 20 mL/min) over eight runs. C9-amino-doxycycline was isolated as its TFA salt as an orange-brown solid (0.592 g, 45.9%). ^1^H NMR (CD_3_OD, 500 MHz) δ 7.58 (d, *J* = 8.3 Hz, 1H), 7.10 (d, *J* = 8.4 Hz, 1H), 4.43 (s, 1H), 3.60 (dd, *J* = 11.4, 8.3 Hz, 1H), 2.96 (br s, 6H), 2.90-2.79 (m, 2H), 2.64 (dd, *J* = 12.3, 8.3 Hz, 1H), 1.58 (d, *J* = 6.9 Hz, 3H). HRMS ESI+ (m/z): 460.1712 (Predicted [M+H]^+^ for C_22_H_25_N_3_O_8_ is 460.1714).

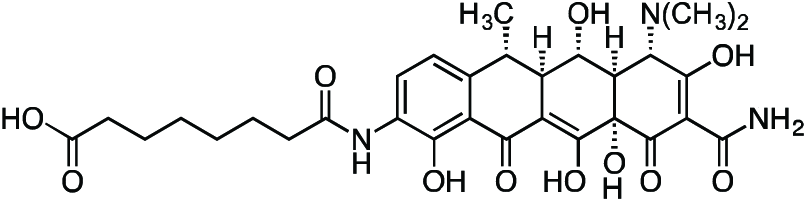

#### Doxycycline SILAC Probe

A flask was charged with C9-amino-doxycycline (0.026 g, 0.045 mmol), HATU (0.018 g, 0.047 mmol), and suberic acid (0.009 g, 0.052 mmol). The reaction components were dissolved with 1 mL DMF, followed by addition of triethylamine (12.5 μL) to give an orange solution. The reaction mixture was then stirred for 90 min at 23 °C. The crude reaction mixture was diluted to 10 mL with 10% CH3CN/H2O + 0.1% TFA, syringe filtered through a 0.2 micron PTFE filter, and purified by prep RP-HPLC on a Waters SunFire Prep-C18, 10 μm, 21.2 x 250 mm column (20 to 40% CH3CN/H2O + 0.1% TFA, 45 min, 20 mL/min). The desired doxycycline SILAC probe carboxylic acid TFA salt was isolated as a yellow solid (15.8 mg, 47.8%). ^1^H NMR (CD_3_OD, 500 MHz) δ 8.15 (d, *J* = 8.3 Hz, 1H), 6.96 (d, *J* = 8.4 Hz, 1H), 4.38 (s, 1H), 3.58 (dd, *J* = 11.4, 8.2 Hz, 1H), 2.94 (s, 6H), 2.85 – 2.72 (m, 2H), 2.59 (dd, *J* = 12.2, 8.3 Hz, 1H), 2.47 (t, *J* = 7.5 Hz, 2H), 2.31 (t, *J* = 7.4 Hz, 2H), 1.73 (p, *J* = 7.1 Hz, 2H), 1.64 (p, *J* = 7.3 Hz, 2H), 1.56 (d, *J* = 6.9 Hz, 3H), 1.45-1.42 (m, 4H). HRMS ESI+ (m/z): 616.2507 (Predicted [M+H]^+^ for C_30_H_37_N_3_O_11_ is 616.2501).

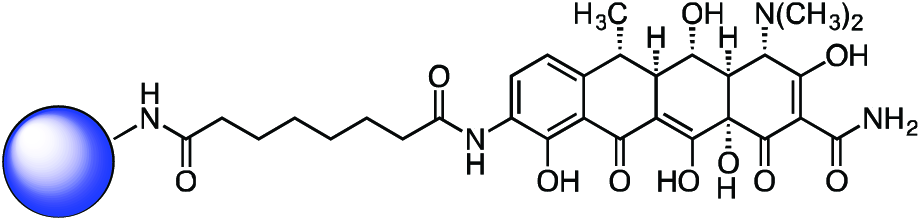

#### Immobilized Doxycycline SILAC Probe (2)

To a solution of the doxycycline SILAC probe carboxylic acid (8.5 mg, 0.012 mmol) and 1-hydroxypyrrolidine-2,5-dione (2.1 mg, 0.018 mmol) in DMSO (800 μl) was added EDC (3.50 mg, 0.018 mmol). After 3h, additional 1-hydroxypyrrolidine-2,5-dione (2.1 mg, 0.018 mmol) and EDC (3.50 mg, 0.018 mmol) were added. After 4h, LC-MS analysis showed 95% activation of the starting material corresponding to 11.4 μmol of activated material.

Affi-Gel 102 beads (12 μmol) were washed with 5 x anhydrous DMSO. The beads were then suspended in anhydrous DMSO (500 μL). The activated solution (1.5 μmol) was added to the bead suspension based on the 12.5% loading level and to this was added triethylamine (8.36 μL, 60.0 μmol). The reaction mixture was shaken at 23 °C for 1 h at which point LC-MS analysis showed complete loss of signal for the activated material. The beads were washed with DMSO and PBS, followed by final suspension in PBS (1.5 mL) for storage of the immobilized doxycycline SILAC probe.

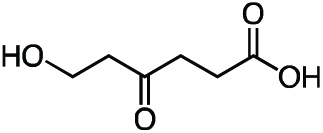

#### 6-Hydroxy-4-oxohexanoic acid

2-Oxepane-1,5-dione(Latere et al., 2002) (1.24 g, 9.68 mmol, 1 equiv) was dissolved in 50 mL dichloromethane and the resulting solution was stirred and cooled to 4 °C. Water (10 mL) was added to the stirring solution to give a biphasic mixture, followed by slow dropwise addition of 10 mL of a saturated potassium carbonate solution. After 30 min, the reaction mixture was warmed to 23 °C and the layers were separated. The aqueous fraction was cooled to 4 °C, followed by slow dropwise addition of an aqueous 10% v/v hydrochloric acid solution until the mixture had reached pH 2.0. The aqueous fraction was then extracted with one 50-mL portion of ethyl acetate. Solid sodium chloride was added to the aqueous fraction until it became saturated, followed by extraction with four 50-mL portions of ethyl acetate. The remaining aqueous fraction was concentrated under reduced pressure, and the resultant crude mixture was triturated with two 100-mL portions of ethyl acetate. The ethyl acetate extracts were combined, dried over magnesium sulfate, filtered, and concentrated to give 6-hydroxy-4-oxohexanoic acid (1.30 g, 92%) as a viscous pale yellow oil. ^1^H NMR (500 MHz, CDCl_3_), δ: 6.42 (br s, 1H), 3.87 (t, *J* = 5.5 Hz, 2H), 2.75 (t, *J* = 6.3 Hz, 2H), 2.72 (t, *J* = 5.5 Hz, 2H), 2.64 (dd, *J* = 7.3, 5.4 Hz, 2H). HRMS ESI-(m/z): 145.0506 (Predicted [M-H]- for C_6_H_10_O_4_ is 145.0506).

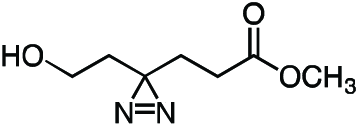

#### Methyl 6-hydroxy-4,4’-azihexanoate

A solution of 7N ammonia in anhydrous methanol (60 mL, 420 mmol, 67.7 equiv) was added to neat 6-hydroxy-4-oxohexanoic acid (0.906 g, 6.20 mmol, 1 equiv) at 4 °C. After complete dissolution of **7**, the solution was cooled to −78 °C in an acetone/dry ice bath and held at that temperature for 5 h. Hydroxylamine *O*-sulfonic acid (0.841 g, 7.44 mmol, 1.2 equiv) in 5 mL anhydrous methanol was added dropwise via cannula at −78 °C, resulting in formation of a white precipitate. After 30 min, the acetone/dry ice bath was removed, and the reaction mixture was gradually warmed to 23 °C and stirred at that temperature for 16 h. The precipitate was removed by filtration, and the resulting filtrate was concentrated. The crude residue was re-suspended in 30 mL methanol, followed by addition of triethylamine (2.60 mL, 18.65 mmol, 3 equiv). The solution was then cooled to 4 °C, and iodine was added in sequential ~200 mg portions until a reddish-brown color persisted. The reaction mixture was then warmed to 23 °C, stirred at that temperature for 1 h, and then concentrated. The crude orange residue was dissolved in 30 mL anhydrous acetonitrile, followed by addition of cesium carbonate (6.10 g, 18.72 mmol, 3 equiv) and methyl iodide (1.6 mL, 25.70 mmol, 4.15 equiv), respectively. The resulting suspension was stirred for 16 h at 23 °C. The reaction mixture was partitioned between 50 mL water and 50 mL dichloromethane. The layers were separated, and the aqueous fraction was extracted with an additional three 50-mL portions of dichloromethane. The combined organic extracts were washed once with 50 mL aqueous saturated brine solution, dried over magnesium sulfate, filtered, and concentrated. The crude residue was purified by flash column chromatography (0 to 100% ethyl acetate/hexanes gradient over 20 column volumes) on an 80 g silica gel column (Teledyne ISCO). Methyl 6-hydroxy-4,4’-azihexanoate (0.398 g, 37.1%) was obtained as of a colorless oil. ^1^H NMR (500 MHz, CDCl_3_), δ: 3.69 (s, 3H), 3.51 (q, *J* = 5.7 Hz, 2H), 2.11 (t, *J* = 7.4 Hz, 2H), 1.98 (br dd, *J* = 12.6, 6.4 Hz, 1H), 1.97 (d, *J* = 8.0 Hz, 2H), 1.83 (t, *J* = 7.4 Hz, 2H), 1.64 (t, *J* = 6.0 Hz, 2H). HRMS ESI+ (m/z): 173.0920 (Predicted [M+H]^+^ for C7H_12_N_2_O3 is 173.0921).

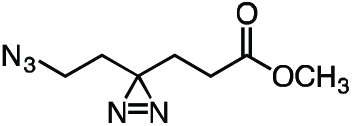

#### Methyl 6-azido-4,4’-azihexanoate

Triethylamine (0.165 mL, 1.184 mmol, 2.04 equiv) was added to a stirring solution of methyl 6-hydroxy-4,4’-azihexanoate (0.100 g, 0.581 mmol, 1 equiv) and *p*-toluenesulfonyl chloride (0.138 g, 0.726 mmol, 1.25 equiv) in 5 mL anhydrous dichloromethane at 4 °C. The resulting clear, colorless solution was stirred at that temperature for 30 min. The reaction mixture was then warmed to 23 °C and held at that temperature for 16 h. The reaction mixture was concentrated under reduced pressure, and the crude residue was re-suspended in 5 mL anhydrous dimethylformamide. Sodium azide (0.113 g, 1.742 mmol, 3 equiv) was added to the reaction mixture in a single portion at 23 °C, and the resulting suspension was stirred for 5 h. The reaction mixture was diluted with 10 mL water and then extracted with three 15-mL portions of diethyl ether. The combined ether extracts were washed once with 15 mL aqueous saturated brine solution, dried over magnesium sulfate, filtered, and concentrated. The crude residue was purified by flash column chromatography (0 to 30% ethyl acetate/hexanes gradient over 20 column volumes) on a 12 g silica gel column (Teledyne ISCO). Methyl 6-azido-4,4’-azihexanoate (0.106 g, 93%) was isolated as of a colorless oil. ^1^H NMR (500 MHz, CDCl_3_), δ: 3.69 (s, 3H), 3.16 (t, *J* = 6.8 Hz, 2H), 2.12 (t, *J* = 7.6 Hz, 2H), 1.80 (t, *J* = 7.5 Hz, 2H), 1.66 (t, *J* = 6.8 Hz, 2H). HRMS ESI+ (m/z): 220.0802 (Predicted [M+Na]^+^ for C7H_11_N5O2 is 220.0805).

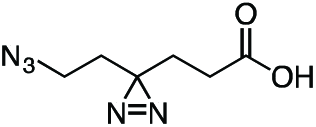

#### 6-Azido-4,4’-azihexanoic acid

Lithium hydroxide (0.121 g in 5 mL water, 5.07 mmol, 10 equiv) was added slowly dropwise to a stirring solution of methyl 6-azido-4,4’-azihexanoate (0.100 g, 0.507 mmol, 1 equiv) in 5 mL THF at 4 °C. After 1 h, the reaction mixture was partitioned between 5 mL dichloromethane and 5 mL water. The layers were separated and the aqueous fraction was cooled to 4 °C, followed by slow dropwise addition of an aqueous 10% v/v hydrochloric acid solution until the mixture had reached pH 2.0. Sodium chloride was then added to the aqueous fraction until it became saturated, followed by extraction with four 15-mL portions of ethyl acetate. The combined ethyl acetate extracts were washed with 10 mL aqueous saturated brine, dried over magnesium sulfate, filtered, and concentrated to provide 6-azido-4,4’-azihexanoic acid (83.6 mg, 90%) as a colorless oil. ^1^H NMR (500 MHz, CDCl_3_), δ: 10.60 (br s, 1H), 3.16 (t, *J* = 6.8 Hz, 2H), 2.17 (t, *J* = 7.6 Hz, 2H), 1.81 (dd, *J* = 7.8, 7.2 Hz, 2H), 1.66 (t, *J* = 6.8 Hz, 2H). HRMS ESI-(m/z): 182.0684 (Predicted [M-H]^−^ for C6H9N5O2 is 182.0683).

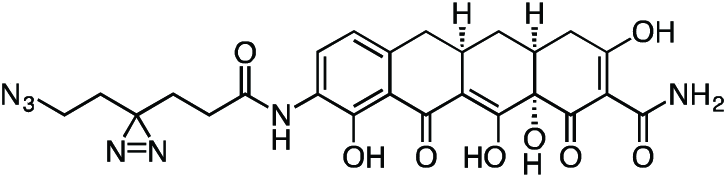

#### Active Col-3 Probe

A flame-dried, round-bottom flask was charged with 9-amino-Col-3 (0.0120 g, 0.031 mmol), HATU (0.0148 g, 0.039 mmol), and 6-azido-4,4’-azihexanoic acid (0.0096 g, 0.052 mmol). The reaction components were dissolved in 0.5 mL anhydrous DMF, followed by addition of triethylamine (15 μL, 0.108 mmol) to give an orange solution. After 3 h, the reaction mixture was diluted with 2 mL acetonitrile + 0.1% TFA and 1.5 mL water + 0.1% TFA, syringe filtered, and purified by prep RP-HPLC on Waters SunFire C18 (20 to 80% CH_3_CN/H_2_O + 0.1% TFA, 45 min, 20 mL/min). The active Col-3 probe TFA salt was isolated as a yellow solid (8.0 mg, 46.7%). ^1^H NMR (500 MHz, CD_3_OD), δ: 8.05 (d, *J* = 8.0 Hz, 1H), 6.74 (d, *J* = 8.1 Hz, 1H), 3.31 – 3.20 (m, 3H), 2.93 – 2.89 (m, 1H), 2.81 (dd, *J* = 15.3, 4.4 Hz, 1H), 2.56 – 2.44 (m, 2H), 2.40 (d, *J* = 18.4 Hz, 1H), 2.33 (t, *J* = 7.6 Hz, 2H), 2.10 – 2.01 (m, 1H), 1.85 (t, *J* = 7.6 Hz, 2H), 1.71 (t, *J* = 6.8 Hz, 2H), 1.58 (q, *J* = 13.0 Hz, 1H). HRMS ESI+ (m/z): 552.1834 (Predicted [M+H]^+^ for C25H_2_5N_7_O_8_ is 552.1837).

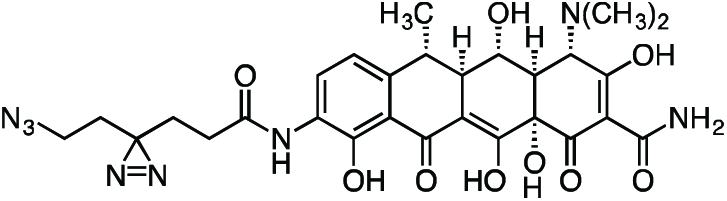

#### Active Doxycycline Probe

A flame-dried, round-bottom flask was charged with 9-amino-doxycycline (0.0495 g, 0.108 mmol), HATU (0.0615 g, 0.162 mmol), and 6-azido-4,4’-azihexanoic acid (0.0237 g, 0.129 mmol). The reaction components were dissolved in 1 mL anhydrous DMF, followed by addition of triethylamine (45 μL, 0.323 mmol) to give an orange solution. The reaction mixture was then stirred for 90 min at 23 °C. The crude reaction mixture was diluted to 10 mL with 10% CH3CN/H2O + 0.1% TFA, syringe filtered through a 0.2 micron PTFE filter, and purified by prep RP-HPLC on a Waters SunFire Prep-C18, 10 μm, 21.2 x 250 mm column (20 to 40% CH3CN/H2O + 0.1% TFA, 45 min, 20 mL/min) over 2 runs. The active doxycycline probe TFA salt was isolated as a bright, yellow solid (26.4 mg, 33.1 %). ^1^H NMR (CD3OD, 500 MHz) δ 8.18 (d, *J* = 8.3 Hz, 1H), 6.92 (d, *J* = 8.5 Hz, 1H), 4.43 (s, 1H), 3.57 (dd, *J*
 = 11.5, 8.3 Hz, 1H), 3.24 (t, *J* = 6.7 Hz, 2H), 2.97 (s, 6H), 2.82 (d, *J* = 11.4, 1.2 Hz, 1H), 2.72 (dq, *J* = 13.5, 6.9 Hz, 1H), 2.55 (dd, *J* = 12.2, 8.3 Hz, 1H), 2.35 (t, *J* = 8.3, 6.8 Hz, 2H), 1.86 (t, *J* = 7.6 Hz, 2H), 1.71 (t, *J* = 6.7 Hz, 2H), 1.54 (d, *J* = 6.8 Hz, 3H). HRMS ESI+ (m/z): 625.2358 (Predicted [M+H]^+^ for for C_2_8H32N_8_O_9_ is 625.2365).

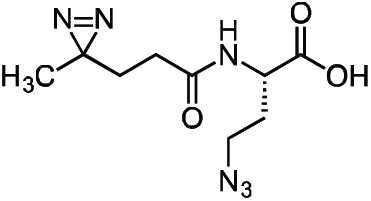

#### Inactive Diazirine-Azide Linker

A flame-dried, round-bottom flask was charged with L-azidohomoalanine (0.030 g, 0.208 mmol) and NHS-diazirine (0.050 g, 0.222 mmol) under argon. The reaction components were dissolved in 2 mL anhydrous acetonitrile, followed by the addition of triethylamine (145 μL, 1.041 mmol). After 1 h, the crude mixture was concentrated under reduced pressure. The reaction mixture was diluted with 5 mL 20% CH3CN/H2O + 0.1% TFA, syringe filtered and purified by prep RP-HPLC on Waters SunFire C18 (20 to 80% CH3CN/H2O + 0.1% TFA, 45 min, 20 mL/min). Isolated the inactive diazirine-azide linker as a colorless oil (40.1 mg, 75.8 %) that solidified upon standing at −20 °C. ^1^H NMR (500 MHz, CDCl_3_), δ: 6.74 (s, 1H), 4.48 – 4.35 (m, 1H), 3.47 – 3.35 (m, 2H), 2.12 – 2.02 (m, 3H), 1.91 −1.83 (m, 1H), 1.67 – 1.59 (m, 2H), 1.01 – 0.99 (m, 3H). HRMS ESI-(m/z): 253.1057 (Predicted [M-H]- for C9H_14_N6O3 is 253.1055).

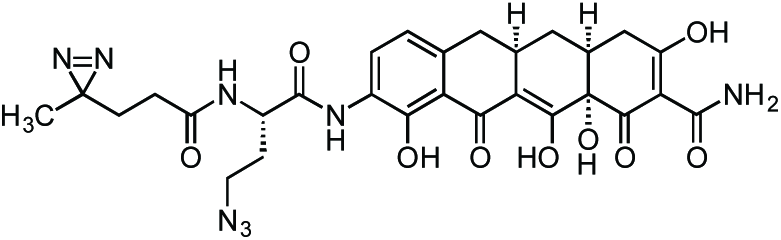

#### Inactive Col-3 Probe

A flame-dried, round-bottom flask was charged with 9-amino-Col-3 (0.0125 g, 0.032 mmol), HATU (0.0140 g, 0.037 mmol), and 6-azido-4,4’-azihexanoic acid (0.009 g, 0.035 mmol). The reaction components were dissolved in 1 mL anhydrous DMF, followed by addition of triethylamine (14 μL, 0.097 mmol) to give an orange solution. The reaction mixture was then stirred for 90 min at 23 °C. The crude reaction mixture was diluted to 10 mL with 10% CH3CN/H2O + 0.1% TFA, syringe filtered through a 0.2 micron PTFE filter, and purified by prep RP-HPLC on a Waters SunFire Prep-C18, 10 μm, 21.2 x 250 mm column (20 to 40% CH3CN/H2O + 0.1% TFA, 45 min, 20 mL/min). The active Col-3 probe TFA salt was isolated as a pale, yellow solid (13.1 mg, 65.0 %). ^1^H NMR (500 MHz, CD_3_OD), δ: 8.07 (dd, *J* = 8.1, 3.0 Hz, 1H), 6.70 (d, *J* = 8.1 Hz, 1H), 4.67 (dd, *J* = 9.1, 5.2 Hz, 1H), 3.54 – 3.42 (m, 2H), 3.24 (dd, *J* = 18.2, 5.6 Hz, 1H), 2.90 – 2.81 (m, 1H), 2.77 (dd, *J* = 15.2, 4.3 Hz, 1H), 2.52 – 2.33 (m, 3H), 2.24 – 2.11 (m, 3H), 2.05 – 1.89 (m, 2H), 1.72 (t, *J* = 7.6 Hz, 2H), 1.54 (q, *J* = 12.7 Hz, 1H), 1.01 (s, 3H). HRMS ESI+ (m/z): 623.2203 (Predicted [M+H]^+^ for for C28H30N8O9 is 623.2209).

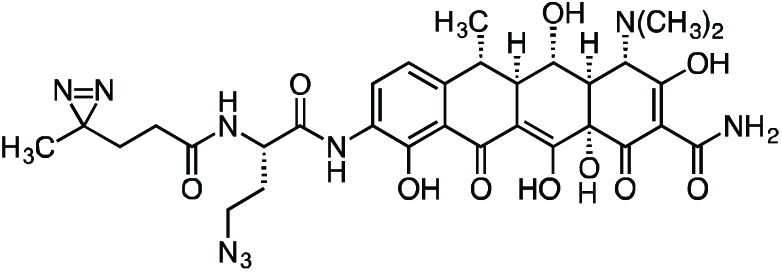

#### Inactive Doxycycline Probe

A flame-dried round-bottom flask was charged with 9-amino-doxycycline (0.0544 g, 0.118 mmol), HATU (0.050 g, 0.131 mmol), and (S)-4-azido-2-(4,4’-azi)butanamido)butanoic acid (0.0327 g, 0.129 mmol). The reaction components were dissolved with 1 mL anhydrous DMF, followed by addition of triethylamine (50 μL, 0.359 mmol) to give an orange solution. The reaction mixture was then stirred for 3 h at 23 °C. The crude reaction syringe filtered through a 0.2 micron PTFE filter, and purified by prep RP-HPLC on a Waters SunFire Prep-C18, 10 μm, 21.2 x 250 mm column (5 to 40% CH3CN/H2O + 0.1% formic acid, 45 min, 15 mL/min). The inactive doxycycline probe HCOOH salt was isolated as a yellow solid (28.7 mg, 34.8%). ^1^H NMR (500 MHz, CD_3_OD) δ: 8.22 (dd, *J* = 8.4, 4.1 Hz, 1H), 6.97 (d, *J* = 8.5 Hz, 1H), 4.69 (dd, *J* = 9.1, 5.3 Hz, 1H), 4.42 (s, 1H), 3.61 – 3.45 (m, 3H), 2.97 (br s, 6H), 2.87 – 2.74 (m, 2H), 2.59 (dd, *J* = 12.2, 8.3 Hz, 1H), 2.26 – 2.15 (m, 3H), 2.05 – 1.93 (m, 1H), 1.74 (td, *J* = 7.7, 2.4 Hz, 2H), 1.56 (d, *J* = 6.9 Hz, 3H), 1.04 (s, 3H). HRMS ESI+ (m/z): 696.2728 (Predicted [M+H]^+^ for for C31H37N9O10 is 696.2736).

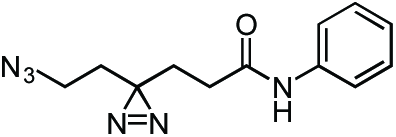

#### Aniline Control Probe

A flame-dried round-bottom flask was charged with HATU, aniline hydrochloride, and 6-azido-4,4’-azihexanoic acid. The reaction components were dissolved in 1 mL anhydrous DMF, followed by addition of triethylamine (50 μL, 0.359 mmol) to give a colorless solution. The reaction mixture was then stirred for 2 h at 23 °C. The reaction mixture was diluted with 2 mL water and extracted with 3 x 5 mL diethyl ether. The combined ether fraction was washed with 5 mL brine, dried over magnesium sulfate, filtered, and concentrated. The crude residue was purified by flash chromatography (0 to 50% ethyl acetate/hexanes gradient over 30 column volumes) on a 4 g silica gel column (Teledyne Isco). The aniline control probe was isolated as a colorless oil (0.022 g, 78.9%). ^1^H NMR (500 MHz, CDCl_3_), δ: 7.51 (d, *J* = 7.5 Hz, 2H), 7.34 (t, *J* = 7.9 Hz, 2H), 7.23 (s, 1H), 7.13 (t, *J* = 7.5 Hz, 1H), 3.19 (t, *J* = 6.7 Hz, 2H), 2.14 (t, *J* = 7.4 Hz, 2H), 1.95 (t, *J* = 7.4 Hz, 2H), 1.71 (t, *J* = 6.7 Hz, 2H). HRMS (ESI): 259.1298 (Predicted [M+H]^+^ for for C1_2_H_14_N_6_O is 259.1302).

### SILAC Affinity Isolation and LC-MS/MS Analysis

Separate cultures of HeLaS3 tetR-GFP cells SILAC-labeled either with L-arginine and L-lysine (light) or L-arginine-^13^C_6_ and L-lysine-^13^C_6_-^15^N_2_ (heavy) were lysed in ice-chilled ModRIPA buffer that was made in RNase free water, containing 1% NP-40, 0.1% sodium deoxycholate, 150 mM NaCl, 0.1 mM DTT, 5 mM MgCl_2_, 50 mM Tris, pH 7.5, 3 μl/ml Protector RNase inhibitor (Roche Applied Science) and EDTA-free protease inhibitors (CompleteTM tablets, Roche Applied Science). Lysates were vortexed intermittently while chilled on ice for 10 min and clarified by centrifugation at 14,000 x g. Protein concentrations of light and heavy lysates were estimated with the Protein Assay Dye Reagent Concentrate (Biorad) and equalized. The protein concentrations of lysates was 1 mg/mL, and affinity enrichments were performed in lysate volumes of 1.4 mL in a 1.5 mL microcentrifuge tube.

Col-3 (in DMSO) at 50-fold excess over the amount of Col-3 on beads was added to 1 mg of light lysate. An equal volume of DMSO was then added to 1 mg of heavy lysate as a control. Both were pre-incubated for 30 minutes. Twenty-five microliters of a 50% slurry in phosphate buffered saline (PBS) of 12.5% loaded Col-3-bead was added to both light and heavy lysates.

Doxycycline (in DMSO) at 100-fold excess over the amount of doxycycline on beads was added to 1 mg of light lysate. An equal volume of DMSO was then added to 1 mg of heavy lysate as a control. Both were pre-incubated for 30 minutes. Twenty-five microliters of a 50% slurry in phosphate buffered saline (PBS) of 12.5% loaded doxycycline was added to both light and heavy lysates. Affinity enrichments were incubated for 3.5 hours on an end-over-end rotator at 4 °C. Following incubation, the beads were pelleted by centrifugation at 1000 x *g*. The supernatant was aspirated, taking care to avoid disturbing the beads. Each tube in a set was washed with ModRIPA buffer twice to remove excess soluble small molecule competitor. Beads from the two tubes were then combined for an extra washing step in ModRIPA. After the third and final wash, beads were collected by centrifugation at 1000 x g, and the wash buffer was aspirated, leaving approximately 20 μL of buffer in the tube.

Proteins enriched in SILAC affinity pulldowns were reduced and alkylated, on bead, in 2 mM DTT and 10 mM iodoacetamide respectively. One part LDS buffer (Invitrogen) was added to three parts sample (including beads), and the tubes were heated to 70 °C for 10 minutes. Proteins were resolved on a 4-12% gradient 1.5 mm thick Bis-Tris gel with MES running buffer (Invitrogen) and Coomassie stained (Simply Blue, Invitrogen). Gel lanes were excised into six pieces and then further cut into 1.5 mm cubes. The gel pieces were further destained in a solution containing 50% EtOH and 50% 50 mM ammonium bicarbonate, then dehydrated in 100% EtOH before addition of sufficient trypsin (12.5 ng/μL) to swell the gel pieces completely. An additional 100 μL of 50 mM ammonium bicarbonate was added before incubating at 37°C overnight on an Eppendorf Thermomixer. Enzymatic digestion was stopped by the addition of 100 μL of 1% TFA to tubes. A second extraction with 300 μL of 0.1% TFA was combined with the first extract and the peptides from each gel slice cleaned up on C18 StageTips (Rappsilber et al., 2007). Peptides were eluted in 50 μL of 80% acetonitrile/0.1% TFA and dried down in an evaporative centrifuge to remove organic solvents. The peptides were then resuspended by vortexing in 7 μL of 0.1% TFA and analyzed by nanoflow-LCMS with an Agilent 1100 with autosampler and an LTQ-Orbitrap.

Peptides were resolved on a 10 cm column, made in-house by packing a self pulled 75 μm I.D. capillary, 15 μm tip (P-2000 laser based puller, Sutter Instruments) column with 3 μm Reprosil-C18-AQ beads (Dr. Maisch GmbH, Ammerbuch-Entringen, Germany) with an analytical flow rate of 200 nL/min and a 58 min linear gradient (~ 0.57% B/min) from 0.1% formic acid in water to 0.1% formic acid/90% acetonitrile. The run time was 50 min for a single sample, including sample loading and column reconditioning. We used a MS method with a master Orbitrap full scan (60,000 resolution) and data dependent LTQ MS/MS scans for the top five precursors (excluding z=1) from the Orbitrap scan. Each cycle was approximately 2 secs long. MS raw files were processed for protein identification and quantitation using MaxQuant v 1.1.1.25, an open source academic software MSQuant (CEBI, open-source http://maxquant.org) IPI human ver.3.68 (http://ebi.ac.uk) with a concatenated decoy database containing randomized sequences from the same database. Common contaminants like bovine serum albumin, trypsin, etc. were also added to the database. Variable modifications used were oxidized methionine, arginine-^13^C_6_, lysine-^13^C_6_ ^15^N_2_, and carbamidomethyl-cysteine was used as a fixed modification. The precursor mass tolerance used in the search was 20 ppm for the first search (used for nonlinear mass re-calibration) and 6 p.p.m. for the main search; fragment mass tolerance was 0.5 Da. Only proteins with a minimum of two quantifiable peptides were included in our dataset.

To model log_2_ protein ratio values, we used normal mixture modeling using the Mclust package in R to determine the null distribution. After normalizing and centering on the null distribution the duplicate experiments where analyzed using the Limma package in the R environment to calculate moderated *t*-test *p*, as described previously (Udeshi et al., 2013). To analyze proteins in common in the Col-3 and doxycycline experiments we combined the datasets using Perseus, an open source academic software MSQuant (CEBI, open-source http://maxquant.org), by matching IPI numbers. In order to determine statistical significance for proteins identified in both experiments the ratios for each experimental replicate were standardized by substracting the mean ratio of the replicate and dividing by the standard deviation of the replicate. The four replicates were then treated as replicates and we used the Limma package in the R environment to calculate the moderated *t*-test *p*.

The original mass spectra have been deposited in the public proteomics repository MassIVE and are accessible at ftp://MSV000012345@massive.ucsd.edu when providing the dataset password: mOVERz. If requested, also provide the username: MSV000012345. This data will be made public upon acceptance of the manuscript.

### In cell “click”-selective crosslinking with RNA sequence profiling

A375 cells (1.0 x 10^6^ cells) were plated into 15 cm dishes in 20 mL complete culture media and grown to 80% confluency at 37 °C. Complete culture medium: DMEM + 10% FBS + 100 μg/mL Normocin.

The cells were treated in duplicate with either DMSO (vehicle), 20 μm active Col-3 Probe, 20 μm active Col-3 Probe + 100 μm soluble Col-3, 20 μm Inactive Col-3 Probe, 20 μm active Doxy Probe, 20 μm active Doxy Probe + 100 μm soluble Doxy, 20 μm Inactive Doxy Probe, and 20 μm Aniline control probe in DMEM + 10% FBS, final DMSO concentration 1% v/v. The dishes were incubated for 3 h at 37 °C. Additional duplicate plates of A375 cells were treated with DMSO to be used as non-crosslinked controls.

Following treatment, the cells were washed with 2 x 10 mL ice-cold 1X HBSS at 4 °C on ice. The HBSS was removed completely, and the dishes (except non-crosslinking controls) were subjected to crosslinking at 365 nm for 20 min at 4 °C on ice in a Stratalinker 1800 at 5 cm distance from the UV source. The non-crosslinking controls were kept at 4 °C on ice under ambient light for 20 min. After crosslinking, the cells were immediately lysed in the dishes with 5 mL Trizol reagent, and the RNA was isolated with the Direct-zol RNA Miniprep Kit (Zymo Research). RNAs (40 μg/sample) were conjugated to DBCO-PEG4-Desthiobiotin (Kerafast) via a copper-free click reaction by incubating with 40 U SUPERaseIN and 1 mM DBCO-PEG4-DesthioBiotin (100 μL reactions; 1% v/v DMSO) in water. The samples were incubated at 37 °C in an Eppendorf Thermomixer at 800 rpm for 4 h. After incubation, the samples were purified with the RNA Clean & Concentrator – 25 Kit (Zymo Research), eluting twice in 25 μL RNase-free water.

The RNAs were lyophilized and re-suspended in 18 μL of RNase-free water. The RNAs were transferred to PCR strip tubes and denatured at 95 °C for 60 s, followed by addition of 2 μL RNA Fragmentation Reagent (Ambion) and heating for an additional 90 s. Reactions were immediately stopped by addition of 2 μL Fragmentation Stop Solution, and the samples were placed on ice. RNA samples were purified with the Zymo RNA Clean & Concentrator – 25 Kit, eluting twice in 25 μL RNase-free water. The isolated RNAs were each split into four aliquots and stored frozen at −80 °C.

One aliquot of each RNA sample (~10 μg) was lyophilized and re-suspended in 10 μL end repair mix: 1 μL 10X PNK buffer, 1 μL SUPERaseIN, 1 μL FastAP alkaline phosphatase (Thermo), 2 μL T4 PNK, 5 μL RNase-free water. End repair was carried out at 37 °C for 1 h. After end repair, adapter ligation was directly carried out by addition of 10 μL adapter ligation mix: 1 μL 50 μm adenylated 3’ adapter (/5rApp/AGATCGGAAGAGCGGTTCAG/3ddC/ for probe-treated samples or /5rApp/AGATCGGAAGAGCGGTTCAG/3TEG-DesthioBiotin/ for DMSO-treated samples), 1 μL T4 RNA Ligase Buffer, 1 μL T4 RNA Ligase 1, High Concentration (NEB), 1 μL 100 mM DTT, and 6 μL 50% PEG8000. Samples were incubated for 3 h at 25 °C.

The adapter ligation reaction mixtures were diluted with 10 μL RNAse-free water and purified with the Zymo RNA Clean and Concentrator – 25 kit, eluting twice with 25 μL RNase-free water. Each of the adapter-ligated RNA mixtures was incubated with 1 μL RPPH, 1 μL RecJ_f_, 2 μL NEB Buffer 2, and 6 μL RNase-free water to digest unligated adapter. Digestion was carried out at 37 °C for 1 h. The RNAs were diluted with 40 μL water and purified with the Zymo RNA Clean and Concentrator – 5 kit, eluting twice with 6 μL RNase-free (recovery ~11 μL).

The ligated samples were annealed to RT primer by addition of 2 μL RT primer (4 pmol; /5phos/DDDNNAACCNNNNAGTCGGAAGAGCGTCGTGAT/iSp18/GGATCC/iSp18/TACTGAA CCGC), 1 μL RNasin Plus (Promega), 2 μL 10 mM dNTPs, and 12 μL RNase-free water. The samples were heat denature at 65 °C for 5 min, and then cooled to 25 °C for 1 min (ramp at 2 °C/s). After annealing, 2 μL DTT, 8 μL SSIV Buffer, and 2 μL SuperScript IV (Thermo Life Technologies) were added. Extension was carried out at 25 °C for 3 min, 42 °C for 5 min, and 52 °C for 30 min, followed by cooling to 4 °C. Following reverse transcription, 5 U RNase I_f_ (NEB) was added to each sample followed by addition of 20 μL of Dynabeads MyOne Streptavidin C1 (pre-washed with and re-suspended in Bead Binding Buffer: 100 mM Tris, pH 7.0, 1 M NaCl, 10 mM EDTA, 0.2% Tween20). The streptavidin capture was carried out at 23 °C for 1 h with constant end over end mixing.

Following capture, the beads were placed on a magnet and the supernatant was removed. The beads were then washed with 5 x 500 μL of 4 M Wash Buffer (100 mM Tris, pH 7.0, 4 M NaCl, 10 mM EDTA, 0.2% Tween20) and 2 x 500 μL of 1X PBS.

Desthiobiotin-tagged samples were eluted from the beads with with 2 x 50 μL of elution buffer: 5 mM D-biotin in 1X PBS. Sample elutions were carried out at 37 °C for 30 min at 1500 rpm on an Eppendorf Thermomixer, followed by placing the sample tubes on a magnet and collection of the supernatant.

The eluted cDNAs were purified with the Zymo DNA Clean and Concentrator-5 Kit using the protocol for isolating ssDNA and recovered from the columns over two 8.5 μL elutions in Zymo DNA Elution Buffer (recovery ~16 μL).

Circularization of the cDNAs was then carried out by addition of 2 μL 10X CircLigase II Buffer, 1 μL 50 mM MnCl_2_, and 1 μL CircLigase II. The reactions were incubated at 60 °C for 2 h, and the cDNA was then cleaned up with the Zymo DNA Clean and Concentrator-5 Kit using the protocol for isolating ssDNA. The cDNAs were recovered from the columns over two 10 μL elutions in Zymo DNA Elution Buffer.

Initial PCR amplification (50 μL reactions) was carried out with 25 μL Q5 HotStart Master Mix (NEB), 2 μL water, and 1 μL each of the 10 μm stock primers:

P3_short primer (CTGAACCGCTCTTCCGATCT)

P5_short primer (ACACGACGCTCTTCCGATCT).

Total PCR reaction volumes were 50 μL. A single cycle of amplification was carried out with the following program:

98 °C for 60 s

98 °C for 15 s

62 °C for 30 s

72 °C for 45 s

After PCR amplification, the cDNAs were again cleaned up with the Zymo DNA Clean and Concentrator-5 Kit with two 10 μL elutions with Zymo DNA Elution Buffer.

Amplification and barcoding was carried out with the Universal Illumina Forward Primer and barcoded reverse primers on a qPCR thermal cycler. Amplifications were monitored by addition of 0.25x Sybr Green and passive 1X ROX dye (Thermo Fisher). Libraries of known concentration (4 nM) were amplified in parallel to identify the optimal number of cycles to achieve a final concentration of ~4-20 nM.

Universal Forward Primer: 5′-AATGATACGGCGACCACCGAGATCTACACTCTTTCCCTACACGACGCTCTTCCGATC*T-3′

Barcoded Reverse Primers: 5′-CAAGCAGAAGACGGCATACGAGATNNNNNNGTGACTGGAGTTCACTGAACCGCTCTTCCGATCT-3′

NNNNNN = 6 bp barcode

Total PCR reaction volumes were 50 μL. Total of 4–10 cycles of amplification was carried out with the following program:

98 °C for 60 s

1 cycle with

98 °C for 15 s

62 °C for 20 s

72 °C for 60 s

3–9 cycles with

98 °C for 15 s

70 °C for 20 s

72 °C for 60 s

After PCR amplification, the cDNAs were again purified with the Zymo DNA Clean and Concentrator-5 Kit and recovered with two 5 μL Zymo DNA Elution Buffer. The library samples were gel purified on a native 6% TBE gel, running the samples at 160 V for ~30 min. The gels were stained with SybrGold in 1X TBE buffer, washed once with 1X TBE buffer, and visualized under a blue light transilluminator. The libraries (>150 bp) were carefully excised from the gel, followed by crushing of the gel slices by centrifugation (>16,000 x *g* for 5 min) through a 0.7 mL Eppendorf tube that had been pierced with a 20G needle and nested in a 2 mL Eppendorf tube. The DNA was eluted from the crushed gel by suspending in 250 μL nuclease-free water, followed by heating at 70 °C for 10 min while shaking at 1,000 r.p.m. on an Eppendorf Thermomixer. The gel slurry was transferred to a Corning Costar Spin-X centrifuge tube filters, and the DNA eluate was recovered by spin filtration at 6,500 x *g*. A second elution was carried out by re-suspending the crushed gel in a second aliquot of 250 μL nuclease-free water and repeating the above extraction steps. The combined eluates for each sample were concentrated with 0.5 mL Amicon 10K columns and the sample libraries were analyzed on an Agilent 2200 TapeStaton with a D1000 High Sensitivity ScreenTape to check for removal of residual primer dimers and empty (no insert) amplicons.

The purified libraries were normalized to 1 nM, multiplexed appropriately, and sequenced (single end 1 x 100 bp) on an Illumina HiSeq 2500. Reverse transcription stops were identified from alignment of demultiplexed sequencing reads to human ribosomal RNA sequences with scripts developed by Flynn and co-workers as previously described (Flynn et al., 2016). The original raw sequencing data for this paper will be deposited in the gene expression omnibus upon submission of this manuscript.

### OPP Flow Cytometry

A375 cells (0.8 x 10^5^ cells) were plated into 12-well plates (1 mL/well) in complete media and grown to 70% confluency. Complete culture medium: DMEM + 10% FBS + 100 μg/mL Normocin.

The cells were dosed in triplicate for 1 or 3 h timepoints with either media alone, DMSO, 10 μm Col-3, 25 μm Col-3, 10 μm Doxycycline, 25 μm Doxycycline, or 10 μm Cycloheximide in complete culture medium. The final DMSO concentration was 1% v/v. A replicate set of plates was also dosed at the same drug concentrations but also with 200 nM ISRIB.

After the appropriate timepoint, the nascent peptides from translating ribosomes were pulse-labeled by addition of 20 μm *O*-propargyl puromycin for 1 h in complete media. The media only replicates were treated without OPP as negative controls. The media was then aspirated off and the cells were detached with Accumax (Innovative Cell Technologies). The detached cells were diluted with 1 mL 1X PBS (without Ca/Mg) and transferred to 1.7 mL Eppendorf tubes. The cells were pelleted by centrifugation at 1,000 x *g* and washed with 1 mL 1X PBS. The wash buffer was aspirated off, and the cells were re-pelleted by centrifugation at 1,000 x *g*.

The cells were then fixed by addition of 0.5 mL 1% formaldehyde in 1X PBS at 4 °C. The fixation was carried out on ice for 15 min. The cells were collected by centrifugation at 1,000 x *g* and washed with 1 mL 1X PBS. The wash buffer was aspirated off, and the cells were re-pelleted by centrifugation at 1,000 x *g*.

Cells were permeabilized by re-suspension in 0.2 mL 1X PBS containing 0.1% sapnonin and 3% FBS and incubation for 5 min at 23 °C. The cells were pelleted by centrifugation at 1,000 x *g*, and the permeabilization solution was removed. The Click-IT Cell Reaction Kit (Life Technologies; C10269) was used to tag the nascent peptides with the AlexaFluor 488 azide fluorophore (5 μm) for 30 min.

The cells were then collected by centrifugation at 1,000 x *g* and washed twice with 1 mL 1X PBS containing 0.1% sapnonin and 3% FBS. The cells were then re-suspended in 0.5 mL 1X PBS containing 4 μg/mL DAPI.

The cells were then subjected to flow cytometry analysis on a BD LSR II Flow Cytometry, collecting > 5,000 events per sample and collecting data in the FITC and Pacific Blue channels. Flow cytometry data was then analyzed using FlowJo software.

### Western Blotting

Samples were prepared with cleared cellular lysates in RIPA buffer containing protease/phosphatase inhibitor cocktail (Pierce) and normalized with a BCA assay kit. The lysates were diluted with 4X LDS loading buffer (Invitrogen) + 10X NuPAGE sample reducing agent, mixed via vortex, and heat denatured at 70 °C for 5 min. The samples were then loaded (~10-20 μg protein/lane) onto 4-12% Bis-Tris Gels (Invitrogen) and electrophoresis was carried out at constant voltage: 50 V for 30 min then 150 V for ~90 min in either 1X MOPS or 1X MES Running Buffer with NuPAGE antioxidant (Invitrogen). The MW standard for visualization was the Biotinylated Protein Ladder Detection Pack (Cell Signaling Technology). The protein bands were then transferred to a 0.2 nitrocellulose (Cell Signaling Technology) membrane at 1.25 A (constant) per gel-membrane sandwich for 12 min on a semi-dry Pierce Power Blotter with in Pierce 1-Step Transfer Buffer. The membranes were washed for 5 min in 1X TBS and blocked in 2% nonfat milk in 1X TBST for 1 h at 23 °C. The blocking solution was removed, and the primary antibodies were diluted into in 5% BSA in TBST, followed by incubation with the membranes overnight (~18 h) at 4 °C. Following overnight incubation, primary antibodies were removed, and the blots were washed with 3 x 10 mL 1X TBST for 5 min/wash. The blots were then incubated with the appropriate secondary antibody (from Cell Signaling Technologies) in 5% nonfat milk in 1X TBST. The secondary antibody was then removed, and the blots were washed with 3 x 10 mL 1X TBST for 5 min/wash. The blots were then incubated with the appropriate developing reagent (Luminata Classico, Crescendo, or Forte Reagent (Millipore)) for 1 min and the chemiluminescent signal was visualized on an AlphaInnotech ChemiImager. Following visualization, the blots were washed with 1X TBS, stripped with Pierce PLUS Western Stripping Buffer, and re-washed with 1X TBS. Blots were then re-blocked and re-probed. Stripping and re-probing was carried out no more than 5 times on a single blot.

| **Antibody** | **Supplier (Catalog #; RRID)** | **Dilution** |
| --- | --- | --- |
| Anti-GAPDH (D16H11) XP Rabbit mAb | Cell Signaling (5174) | 1:10,000 |
| Anti-EIF2S1 (phospho S51) [E90] Rabbit mAb | Abcam (ab32157) | 1:2,000 |
| Anti-ATF4 (D4B8) Rabbit mAb | Cell Signaling (11815) | 1:1,000 |
| Anti-EIF2α (D7D3) XP Rabbit mAb | Cell Signaling (5324) | 1:2,500 |
| Anti-ATF3 [EP13502] Rabbit mAb | Abcam (ab200655) | 1:1,000 |
| Anti-IRE1α (phospho S724) [EPR5253] Rabbit mAb | Abcam (ab124945) | 1:1,000 |
| Anti-IRE1α (14C10) Rabbit mAb | Cell Signaling (3294) | 1:1,000 |
| Anti-ATF6 (D4Z8V) Rabbit mAb | Cell Signaling (65880) | 1:1,000 |
| Anti-PERK (D11A8) Rabbit mAb | Cell Signaling (5683) | 1:1,000 |
| Anti-β-actin (13E5) Rabbit mAb | Cell Signaling (4970) | 1:5,000 |
| Anti-α-Tubulin (11H10) Rabbit mAb | Cell Signaling (2125) | 1:5,000 |

### Gene expression analysis

RNA was isolated from cells treated under the appropriate conditions by first washing once with 1X Hank’s Balanced Salt Solution (HBSS) and then directly lysing cells with RiboZol RNA extraction reagent (Amresco). Total RNA was purified with the Direct-zol RNA Miniprep Kit (Zymo Research), according to the manufacturer’s protocol. Following isolation, DNase treatment was performed by incubating with 5U Turbo DNase (Ambion) for 30 min at 37 °C. The RNAs were quantitated by Nanodrop and quality was assessed on either an Agilent 2100 Bioanalyzer or TapeStation 2200. All total RNA samples with RNA Integrity Numbers (RIN) > 7 were deemed suitable for downstream applications.

RNA-Seq libraries were prepared with the Illumina TruSeq RNA Library prep kit and analysis was carried out on 1 x 75 bp sequencing reads as previously described (Trapnell et al., 2012). The original raw sequencing data for this paper will be deposited in the gene expression omnibus upon submission of this manuscript.

Nanostring analysis was carried out at the Whitehead Institute Genome Technology Core with a custom codeset for the target genes of interest.

### Structural analysis

Potential binding sites were analyzed in the 3D structure of the human 80S ribosome determined recently by cryo-electron microscopy (Khatter et al., 2015) using the latest, further refined atomic model (Myasnikov et al., 2016). Atomic models of Doxycline and Col3 were built using the tool “Ligand Builder” available in COOT (Emsley et al., 2010). The built atomic models were placed into the human 80S ribosome structure (PDB code: 5LKS) at the vicinity of the respective predicted cross-linking sites. The models were oriented based on the surface complementarity between the ligands and possible binding area located proximally to the cross-linking sites. The possible interacting residues were inspected visually and identified using COOT. Nucleotides from the rRNA or amino acids of ribosomal proteins within a cut-off distance of 10 Å from the main residues that cross-link with the tetracycline ligands were considered as potential interacting residues. We thereby identified key residues, which appear to be exposed in the vicinity of each cross-link site and that would possibly provide opportunities for hydrogen bonds and stacking interactions for the tetracycline derivatives. To avoid misleading conclusions about ligand moieties in contact with the ribosome we did not attempt precise modelling of the ligands considering the limited precision of positioning ligands in the structure because the sites are different from those known in bacterial ribosomes (Brodersen et al., 2000; Pioletti et al., 2001). Figures were prepared using the software Pymol (www.pymol.org).

## AUTHOR CONTRIBUTIONS

J.D.M. designed and executed research experiments and analyzed data for all project areas. M.S. and C.C. designed and executed SILAC-based affinity isolation experiments and sample analysis. M.S. performed analysis and statistical modeling of mass spectrometry data from SILAC-based affinity isolation experiments. J.A.M. and Z.Z. contributed to Western Blotting experiments. J.A.M. contributed to OPP-flow cytometry translation analysis experiments. L.C. contributed to cellular proliferation assays. E.C. contributed to preparation of tetracycline affinity matrices. S.A.C. oversaw SILAC-based proteomics experiments. S.K.N. and B.P.K. performed structure-based binding site analysis. A.G.M oversaw execution of project goals. J.D.M and A.G.M. wrote the manuscript.

## ACKNOWLEDGMENTS

This work was supported by N.I.H. 5R01CA047148-25 (awarded to A.G.M). J.D.M was supported by a fellowship from the American Cancer Society New England Division – Ellison Foundation. S.K.N. and B.P.K. were supported by CNRS, Association pour la Recherche sur le Cancer (ARC), Ligue contre le Cancer and Institut National du Cancer (INCa). The electron microscope facility was supported by the Alsace Region, the Fondation pour la Recherche Médicale (FRM), INSERM, CNRS, ARC, the French Infrastructure for Integrated Structural Biology (FRISBI) ANR-10-INSB-05-01, and Instruct as part of the European Strategy Forum on Research Infrastructures (ESFRI). We thank Professor Brian Liau for his insight and critique of this manuscript.

## SUPPLEMENTAL INFORMATION

Supplemental Information includes four figures and five data files, which can be found in the online version of the paper.

### CONTACT FOR REAGENT AND RESOURCE SHARING

Further information and requests for resources and reagents should be directed to and will be fulfilled by the Lead Contact, Andrew G. Myers

### Supplemental Figures

**Supplemental Figure 1.**
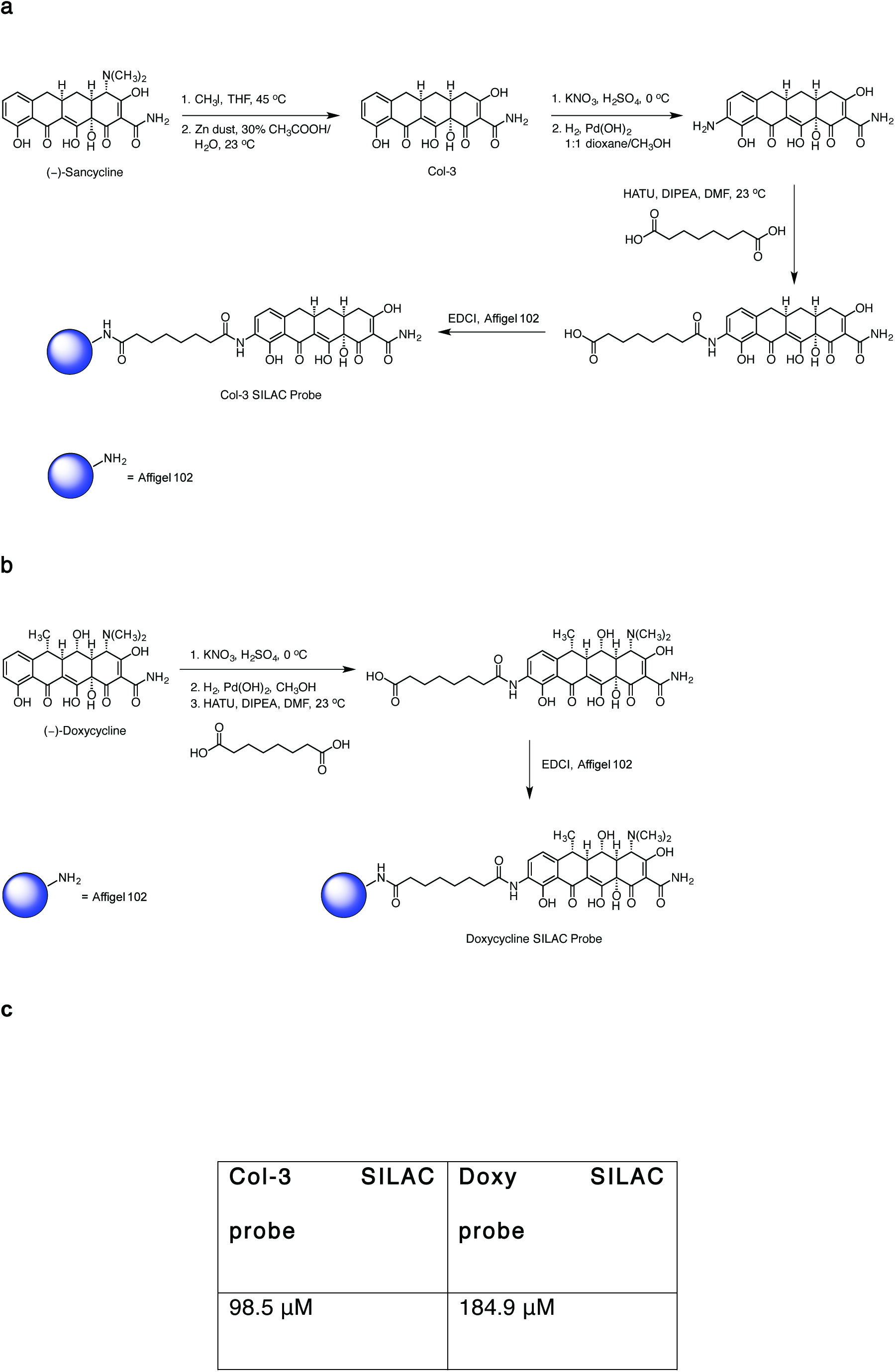
Scheme for tetracycline probe synthesis. **(a)** Synthesis of Col-3 Affinity Probe and **(b)** Synthesis of Doxycycline Affinity Probe and **(c)** GI_50_ values for the tetracycline Col-3 and doxycycline SILAC probes (ethyl amide derivatives)

**Supplemental Figure 2.**
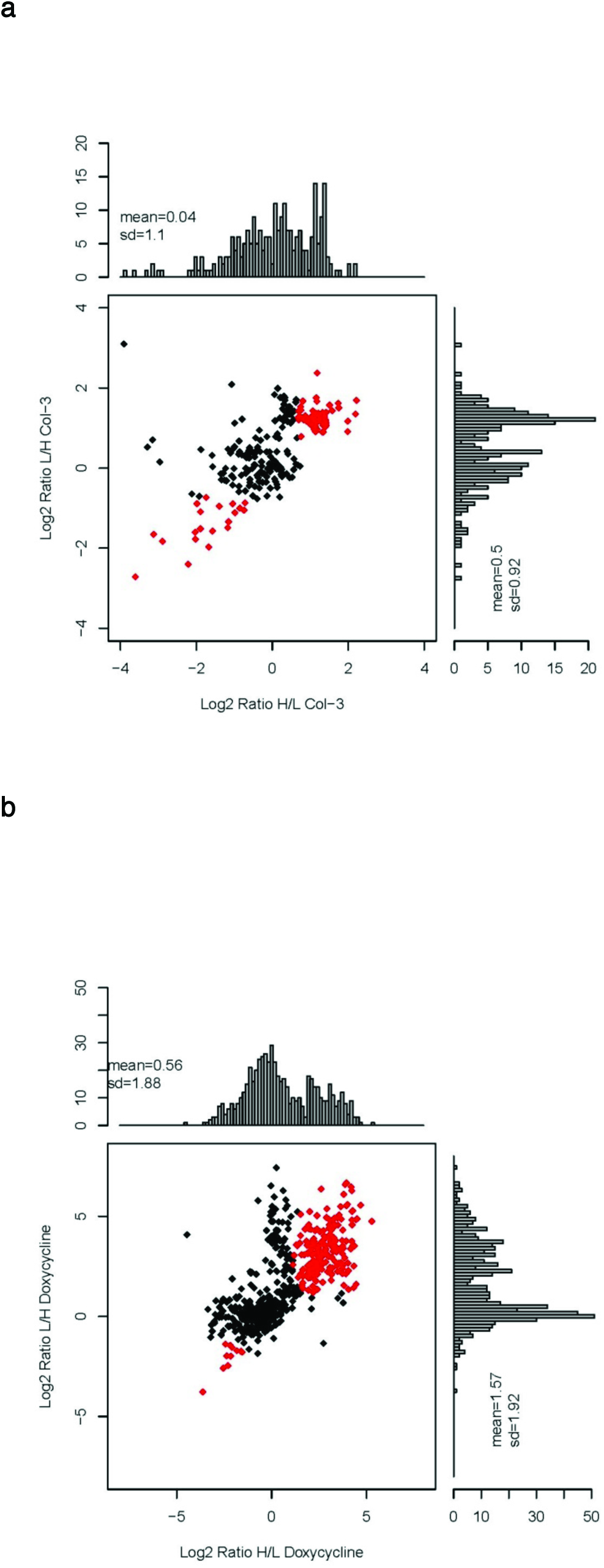
Histograms and scatter plots for proteins identified from SILAC tetracycline affinity isolation experiements. **(a)** Specific proteins identified from Col-3 pulldowns. **(b)** Specific proteins identified from doxycycline pulldown experiments. Proteins with significantly different SILAC ratios (*p* value < 0.05) are shown in red. L = light SILAC lysate fraction and H = heavy SILAC lysate fraction

**Supplemental Figure 3.**
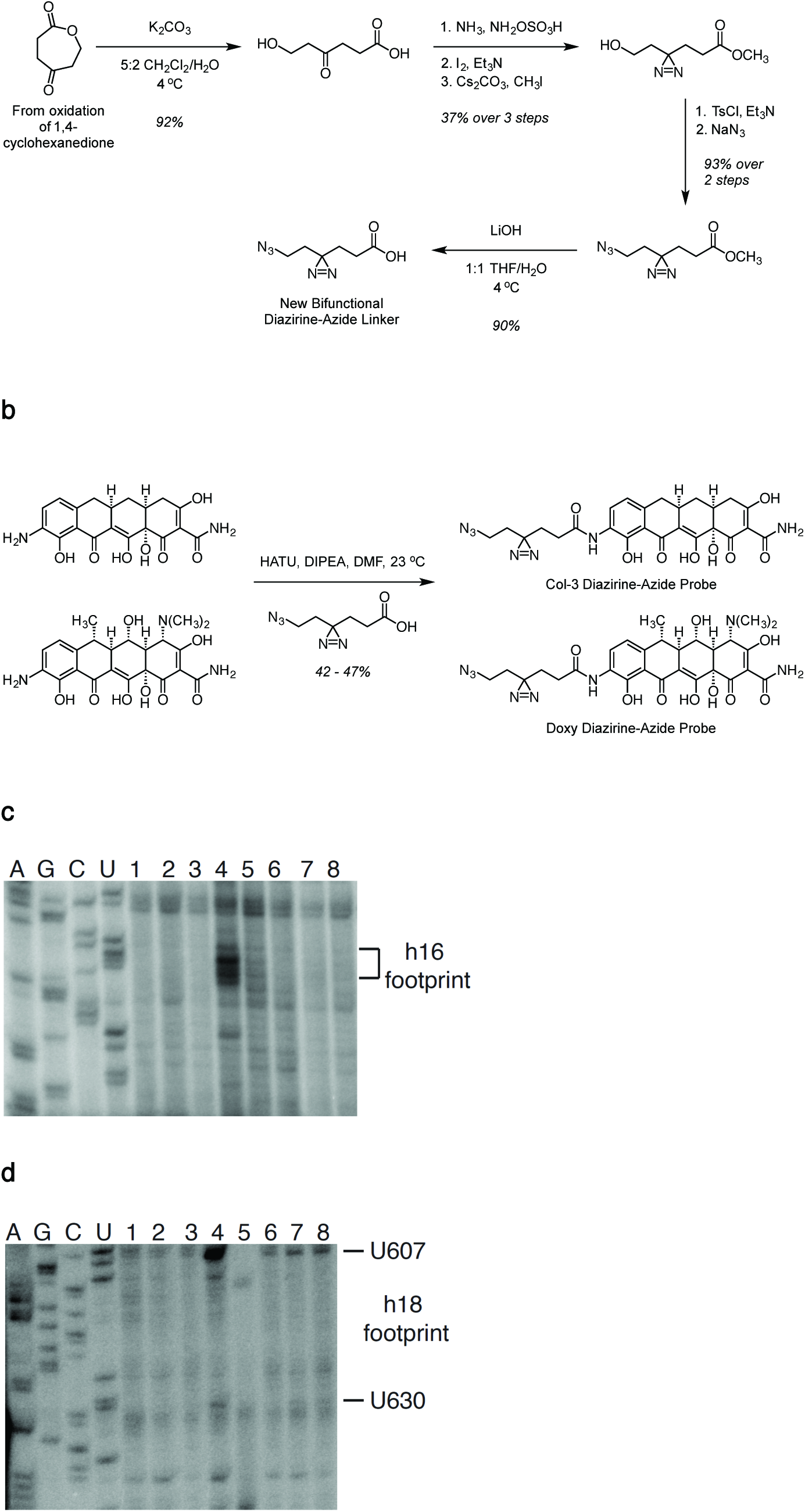
Scheme for the efficient synthesis of bifunctional diazirine-azide tetracycline probes. (**a**) Synthesis of a novel bifunctional diazirine-azide linker. (**b**) Synthesis of bifunctional Col-3 and doxycycline crosslinking probes. (**c**) ^32^P radio-labeled primer extension for footprinting of tetracycline crosslinks on helix h16 (nucleotides 538-542) in A375 cells. (**d**) ^32^P radio-labeled primer extension for footprinting of tetracycline crosslinks on helix h18 (U607 and U630) in A375 cells. Lanes 1-8 correspond to RNA samples from A375 cells treated with DMSO, 20 μm aniline control probe, 20 μm inactive Col-3 control probe, 20 μm Col-3 diazirine-azide probe, 20 μm Col-3 diazirine-azide probe + 100 μm Col-3 soluble competitor, 20 μm inactive doxycycline control probe, 20 μm doxycycline diazirine-azide probe, 20 μm doxycycline diazirine-azide probe + 100 μm doxycycline soluble competitor

**Supplemental Figure 4.**
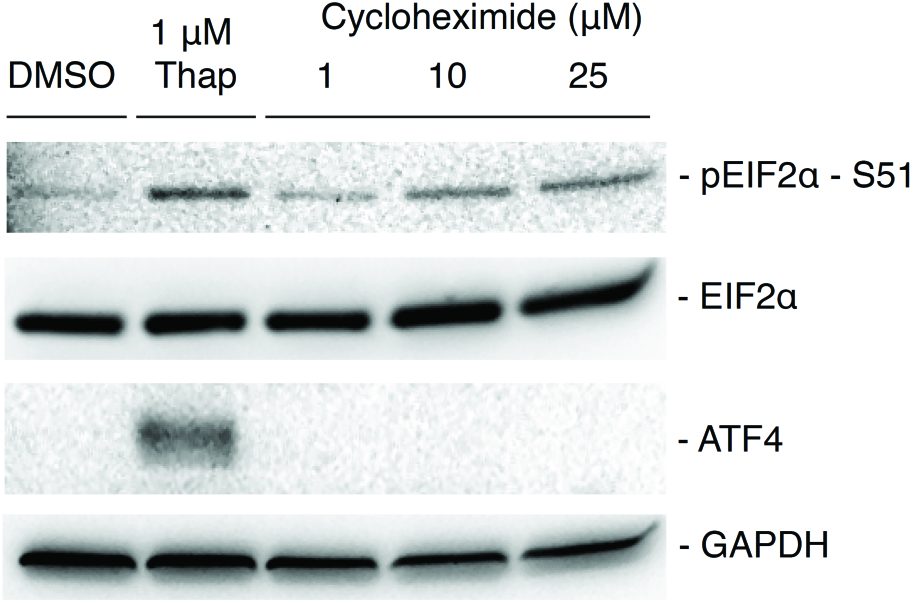
Concentration-dependent phosphorylation of EIF2α in cycloheximide-treated HCT 116 cells at 6 hours.

